# Single nucleus multiomics identifies ZEB1 and MAFB as candidate regulators of Alzheimer’s disease-specific *cis* regulatory elements

**DOI:** 10.1101/2022.10.04.510636

**Authors:** A. G. Anderson, B. B. Rogers, J. M. Loupe, I. Rodriguez-Nunez, S. C. Roberts, L. M. White, J. N. Brazell, W. E. Bunney, B. G. Bunney, S. J. Watson, J. N. Cochran, R. M. Myers, L. F. Rizzardi

## Abstract

Cell type-specific transcriptional differences between brain tissues from donors with Alzheimer’s disease (AD) and unaffected controls have been well-documented, but few studies have rigorously interrogated the regulatory mechanisms responsible for these alterations. We performed single nucleus multiomics (snRNA-seq+snATAC-seq) on 105,332 nuclei isolated from cortical tissues from 7 AD and 8 unaffected donors to identify candidate *cis*-regulatory elements (CREs) involved in AD-associated transcriptional changes. We detected 319,861 significant correlations, or links, between gene expression and cell type-specific transposase accessible regions enriched for active CREs. Among these, 40,831 were unique to AD tissues. Validation experiments confirmed the activity of many regions, including several candidate regulators of *APP* expression. We identified ZEB1 and MAFB as candidate transcription factors playing important roles in AD-specific gene regulation in neurons and microglia, respectively. Microglial links were globally enriched for heritability of AD risk and previously identified active regulatory regions.

## Introduction

Identification of genetic contributors to Alzheimer’s disease has provided critical insights into potential disease mechanisms. Rare, protein-altering variants in *APP, PSEN1*, or *PSEN2* cause early-onset, autosomal dominant AD ^1^, and genome-wide association studies (GWAS) have identified common variants for late-onset AD that increase disease risk to varying degrees ^2–6^. However, the majority of GWAS variants are located in noncoding regions of the genome and many presumably affect gene regulation. Linkage disequilibrium makes identification of the causal variant difficult, particularly for putative regulatory regions where conservation and deleteriousness estimates may not be as informative. Associating common and rare regulatory variants with affected genes is also challenging^7–9^. In addition, disease-associated variants often function only in specific cell types, further complicating interpretation of their effects^10,11^. Thus, determining which genes are contributing to disease requires assessments in specific cell types.

Recent advances in single cell technologies have allowed profiling of gene expression^12–18^ and chromatin accessibility^10^, either separately or in parallel from the same samples^19,20^. While these studies have examined the cell type–specific transcriptional and epigenetic differences between tissues from brain donors with AD and unaffected controls, few have rigorously interrogated the regulatory mechanisms responsible for these alterations^11,21^. Integrating single nucleus RNA-seq (snRNA-seq) and single nucleus ATAC-seq (snATAC-seq) data allows identification of potential *cis* regulatory elements (CREs) by correlating chromatin accessibility with nearby gene expression. Here, we simultaneously measure both gene expression and chromatin accessibility in the same nuclei to identify cell type–specific regulatory regions and their target genes in dorsolateral prefrontal cortex (DLPFC) tissues from both AD and unaffected donors. In addition, we identify regulatory mechanisms unique to nuclei from donors with AD.

## Results

### Cellular diversity within the human dorsolateral prefrontal cortex (DLPFC)

We used the 10X Genomics Multiome technology to perform snATAC-seq and snRNA-seq on nuclei isolated from human postmortem DLPFC tissues from seven individuals diagnosed with AD (mean age 78; Braak stages 4-6) and eight sex-matched unaffected control donors (mean age 63) (**Table S1; Figure 1A**). This assay allows direct mapping of both gene expression and chromatin accessibility within the same nuclei without the need to computationally infer cell type identification during cross-modality integration. After removing low quality nuclei and doublets (*Methods*), we retained a total of 105,332 nuclei with an average of 7,022 nuclei per donor (range of 1,410 - 11,723). We detected a median of 2,659 genes and 11,647 ATAC fragments per cell. We performed normalization and dimensionality reduction for snRNA-seq and snATAC-seq data using Seurat (v4)^22^ and Signac (v1)^23^, respectively. We used weighted-nearest neighbor (WNN) analysis to determine a joint representation of expression and accessibility and identified 36 distinct clusters composed of eight major cell types and their associated subclusters (**Figures 1B, S1A and S1B**). Consistent with previous scRNA-seq data sets^12,13,15,19^, we identified all expected cell types in the brain with similar relative abundances across AD and control donors (**Figures 1B, 1C, and S1C**). Pericytes and endothelial cell clusters contained *<*500 nuclei and were excluded from further analyses. Cluster annotations were supported by both gene expression and promoter accessibility of well-established cell type marker genes (**Figures 1D and 1E**). There were strong correlations in global gene expression across donors within each cell type and between excitatory/inhibitory neurons (**Figure 1F**). The only cell type to display variable correlation values across donors was microglia, a cell type known to be dysregulated in AD. In addition, we identified distinct subpopulations within each major cell type with the exception of oligodendrocyte precursor cells (OPCs), pericytes, and endothelial cells (**Figure S2**). These subtype annotations were consistent with those from prior works^14,22,24^ and distributions were similar across AD and control donors, with the exception of microglia subpopulations and two inhibitory neuron subtypes (Inh 1 and Inh 2; **Figure S2**).

**Figure 1.**
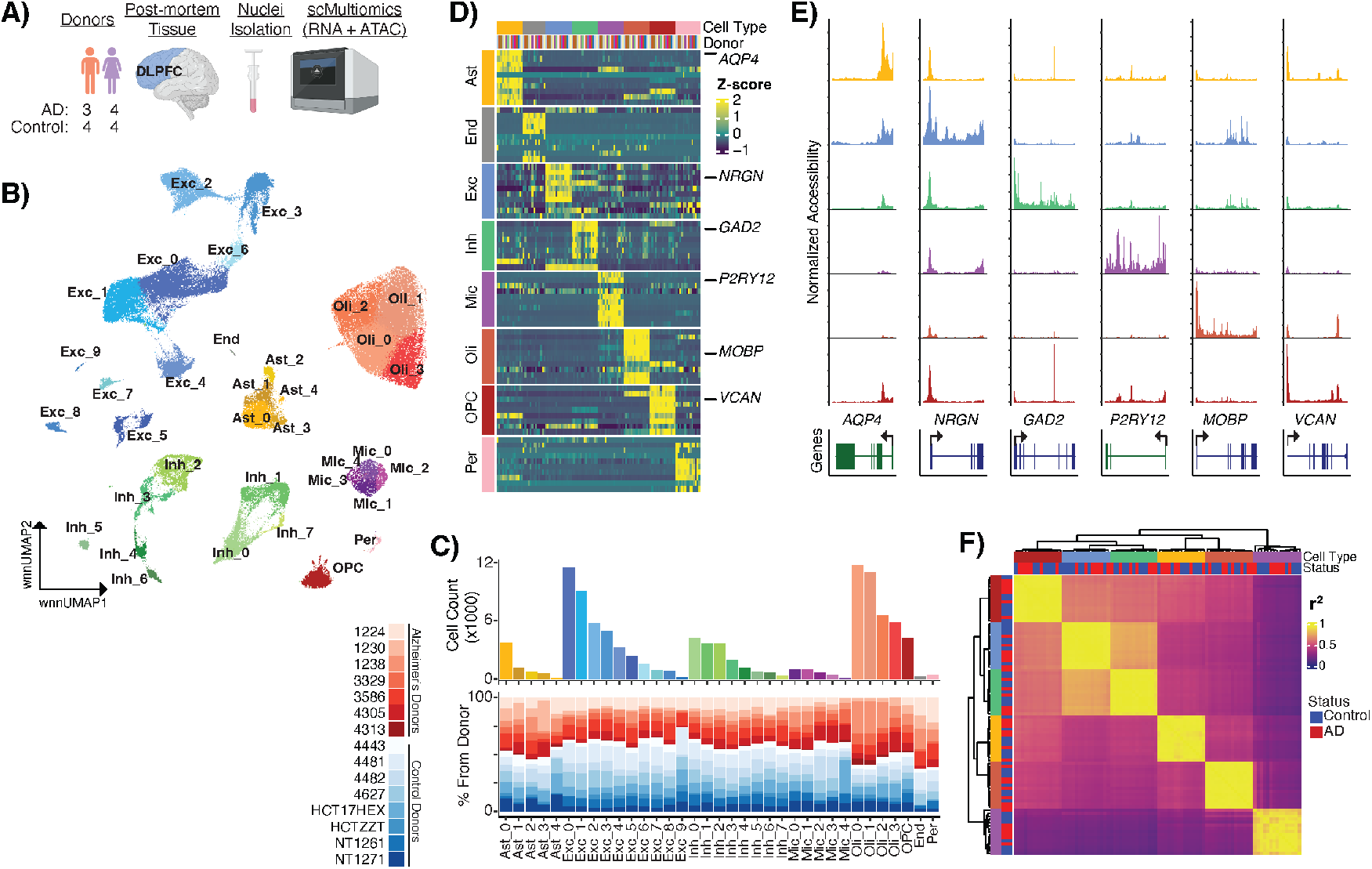
Cellular diversity of DLPFC from Alzheimer’s disease and unaffected donors revealed by single cell multiomics.. **A)** Experimental design. **B)** UMAP visualization of the weighted nearest neighbor (WNN) clustering of single nuclei colored by cell type and cluster assignment. **C)** Total number of cells in each subcluster and the proportion of cells from each individual (AD donors = red; unaffected donors = blue) in the subcluster. **D)** Row-normalized gene expression of scREAD cell type markers. **E)** Chromatin accessibility across cell types for cell type marker genes (indicated below). **F)** Correlation of pseudo-bulked cell type-specific expression profiles between individuals. Colors indicating cell type are consistent throughout the figure.

### Cell type-specific transcriptome changes in Alzheimer’s DLPFC

Within each cell type, we identified differentially expressed genes (DEGs) between AD and control tissues. A total of 911 DEGs were identified after considering sex and age as covariates (**Figure 2A, Table S3**). While significant sex-specific diferences in gene expression between AD and controls have been shown previously^24^, due to our smaller sample size we did not detect such changes. While the majority of DEGs were cell-type specific, 141 were identified across multiple cell types (**Figure 2B**). Of these DEGs, 62 were also identified in both Mathys et al.^12^ and Morabito et al.^19^, including *PTPRG*, which is upregulated in AD microglia across all three studies (**Figures 2C and 2D**). Most DEGs were upregulated in AD and were enriched for cell type–specific gene ontology terms including PDGFR beta signaling in microglia, apoptosis in astrocytes, and Notch and BDNF signaling in oligodendrocytes (**Figure 2E, Table S4**). In contrast, most DEGs downregulated in AD were in neurons and showed enrichment in regulation of tau activity (**Figure 2E, Table S4**).

**Figure 2.**
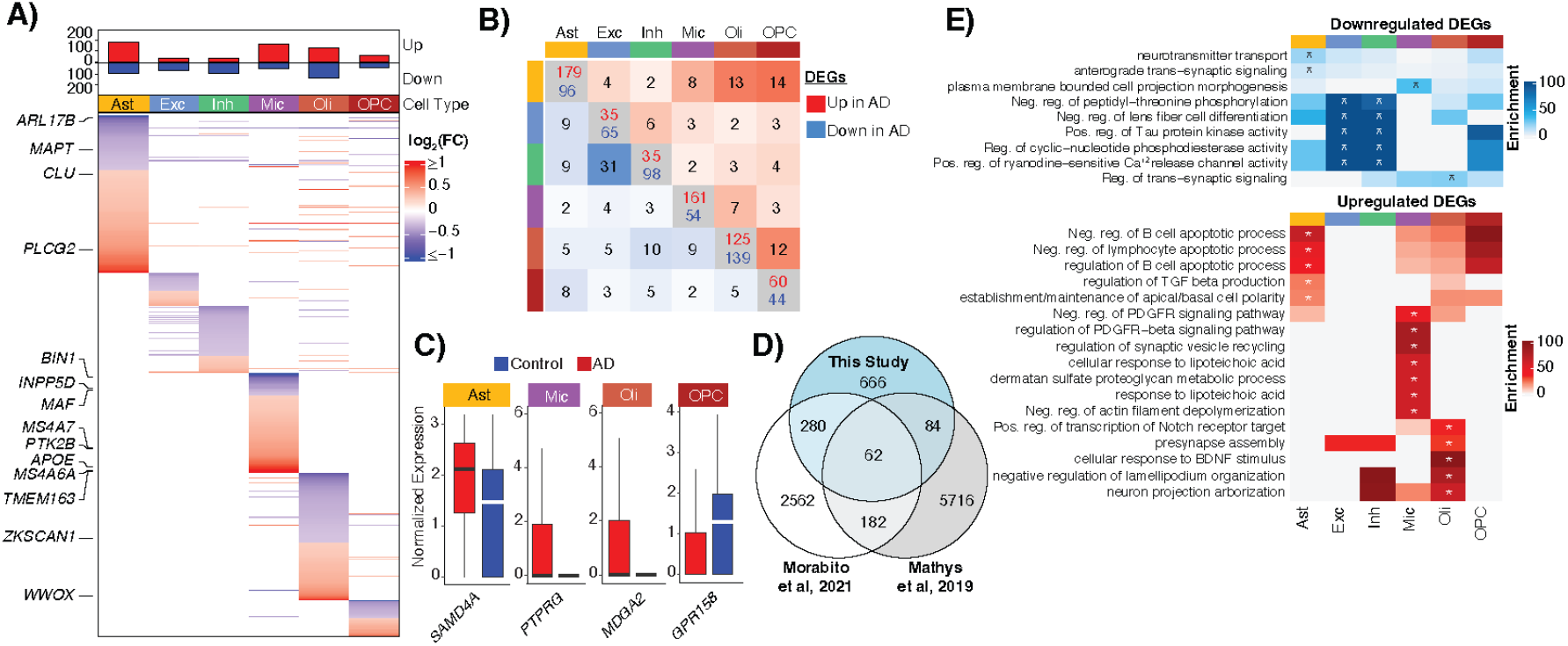
Cell type-specific transcriptome dysregulation in Alzheimer’s DLPFC.. **A)** MAST log(FC) of all up- and down-regulated genes in AD for each cell type. **B)** Number of shared DEGs between cell types in both directions (upper triangle: up-regulated in AD; lower triangle: down-regulated in AD). **C)** Normalized expression of the top DEG in the indicated cell types (log2(FC) *>*1). D) Overlap of DEGs with agreement on cell type and direction with Morabito et al.^19^ and Mathys et al.^12^. **E)** Heatmap showing the odds ratio of the top gene ontology terms for up and down-regulated DEGs within each cell type (* indicates terms with an adjusted p-value *<*0.01).

### Identification of candidate *cis* regulatory elements

Previous single cell studies have characterized altered gene expression in AD brain tissues and cell types^12–14,17,19^, and we observed signals consistent with those studies. Additionally, we sought to leverage single cell multiomics data to identify cell type– and disease-specific CREs and their target genes by correlating gene expression with chromatin accessibility. The cellranger-arc (v2.0) analysis pipeline produces these correlations as “feature linkages”. A feature linkage, or link, is defined as an ATAC peak with a significant correlation, across all nuclei in the data set, between its accessibility and the expression of a linked gene^25^ (**Figure 3A**). We restricted this correlation analysis to consider only peaks within 500 kb of each transcription start site (TSS), as previous studies have found the majority of enhancers are within 50-100 kb of their target genes^26^. We first took the union of ATAC peaks identified in each cell type and only retained those present in ≥2% of cells in at least one cell type for a total of 189,925 peaks. Using this peak set, feature linkages were then calculated independently using gene expression data from either AD or control nuclei allowing classification of linkages as AD-specific, control-specific, or common (*Methods*). Cell type specificity of each link was determined by the cell type(s) in which the ATAC peak was identified. A total of 319,905 peak-gene links were found involving 15,471 linked-genes and 126,213 linked-peaks with a minimum absolute correlation value of 0.2 (**Figure 3A, Table S5**). The median distance between the linked peak and the TSS of the linked gene was 201,506 bp and there was an inverse relationship between absolute correlation value and distance to TSS (**Figure S3A**).

**Figure 3.**
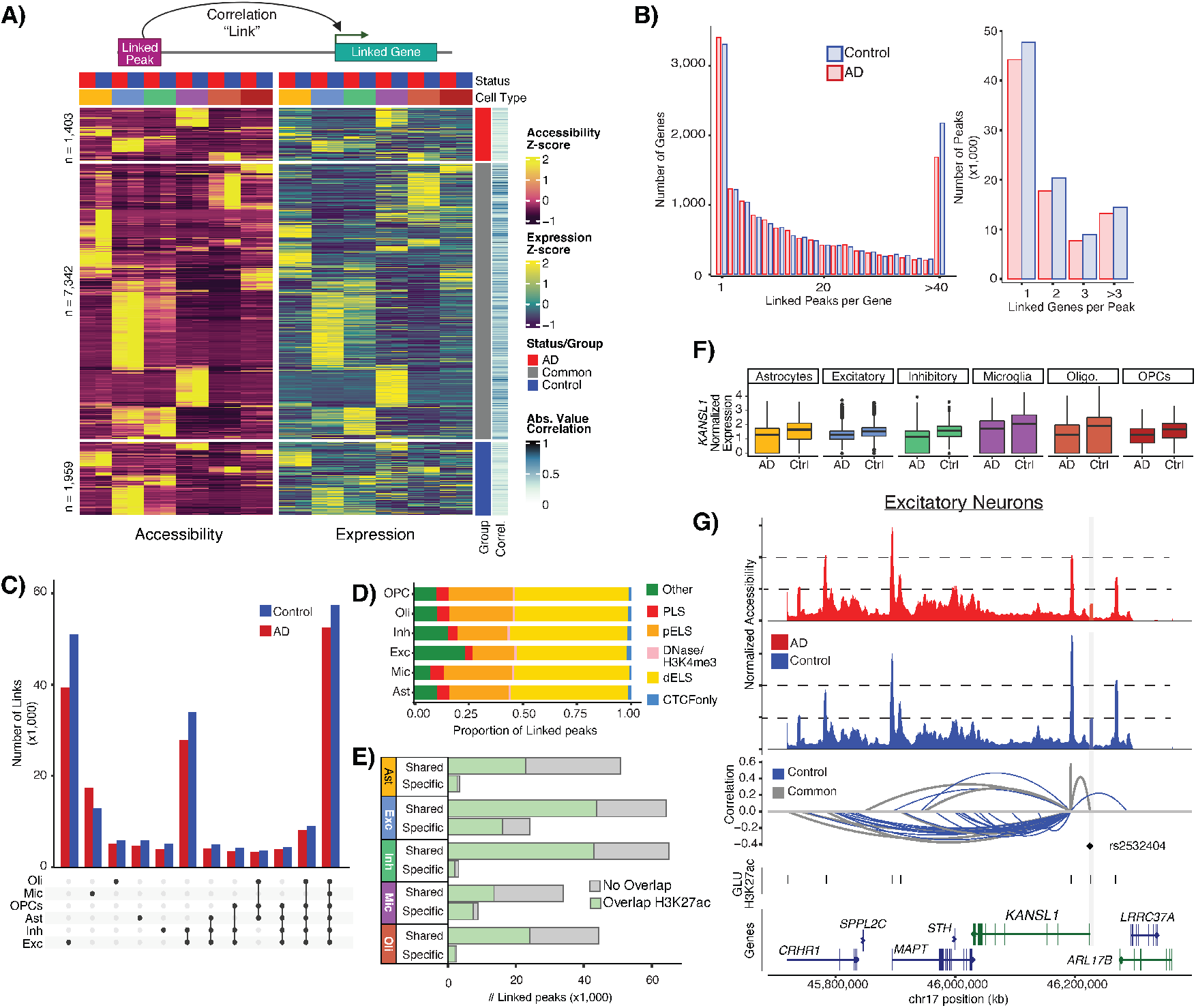
Identification of candidate CREs. **A)** Schematic of gene-peak association (top). Heatmap of row-normalized accessibility and expression for the most correlated gene-peak link for each gene (bottom). Columns are pseudo-bulked on cell type and disease status. **B)** Distributions of the number of linked peaks per gene (left) and the number of linked genes per peak (right) for AD (red) and control (blue) samples. **C)** Total number of links per cell type for AD and control. Cell type of the link is assigned by the cell type in which the peak was called. **D)** ENCODE annotation of linked peaks by cell type. **E)** Shared (across cell types) and cell type-specific linked peaks that overlap H3K27ac of the corresponding cell type. **F)** Normalized expression of *KANSL1* from AD and control samples in each cell type. Expression is significantly different in AD vs control for all cell types. **G)** Linkage plot for all links to *KANSL1*. Top panels: coverage plot of pseudo-bulked accessibility in excitatory neurons separated by status (red = AD; blue = control). Bottom panel: significant AD and control peak-gene links. Arc height represents strength and direction of correlation. Arc color indicates if the link was identified in both AD and control (“common”, gray) or control donors only (blue). A linked peak overlapping a single SNP is highlighted in gray.

For most genes, we identified a similar number of links in both AD (median = 12) and control samples (median = 13). However, we found 1,294 genes had only AD links and 1,596 had only control links (**Figure S3B**). We observed no significant bias when comparing the number of links identified in either AD or control for a given gene (**Figure S3B**). Most genes were linked to multiple peaks across all cell types with a median of 14 linked peaks per gene. However, 16% of genes were linked with 40 or more peaks (**Figure 3B**) and these genes were significantly longer and more highly expressed than those with fewer links (**Figure S3C**).

ATAC peaks often did not interact with only one gene. Nearly 70% (126,213) of the ATAC peaks analyzed were linked to a gene with an average of two genes linked to each peak and a range of 1-21 linked genes (**Figure 3B**). Links ranged from being unique to one cell-type to shared across all.

Almost a third (30.24%) of the links were unique to a single cell type while 21% were common across all cell types (**Figure 3C**). We identified 40,831 AD-specific links and 74,028 control-specific links with the majority of links identified in both (205,046). To evaluate whether linked peaks associate with regulatory regions, we evaluated their overlap with a curated set of candidate CREs identified by ENCODE^27^. We found that linked peaks were significantly enriched for proximal (OR = 1.24, p = 2.4 × 10^−15^) and distal (OR = 1.06, p = 3.06 × 10^−9^) enhancer-like sequences and the proportion of overlap was similar across cell types (**Figure 3D**). As these annotations were not generated in our particular cell types and tissue, we also intersected these linked peaks with regions of H3K27ac previously identified within cell types isolated from prefrontal cortex tissues^11,28^. We found that, on average, 57.5% of linked peaks overlap a H3K27ac peak from the corresponding cell type and this increases to 79% for cell type-specific linked peaks (**Figure 3E**). The majority (76.11%) of linked peaks were positively correlated with gene expression, as is expected given the association between open chromatin and transcriptional activation.

In order to associate DEGs with CREs, a link must be present for that gene. For DEGs identified between AD and control nuclei, 95% had at least one linked peak. Of these DEGs, 69% had a cell type-specific link in the same cell type where the gene was differentially expressed. One example is *KANSL1*, a gene located in the *MAPT* locus that encodes a ubiquitously expressed member of a histone acetyltransferase complex. Loss of function mutations in *KANSL1* result in neurodevelopmental defects and intellectual disability^29^. *KANSL1* was the only DEG identified in all cell types and was downregulated in AD (**Figure 3F**). Nine of the 37 *KANSL1* linked peaks are found in both AD and control donors, and 14 are neuron-specific (**Figure 3G**). One of these linked peaks found in the promoter overlaps an eQTL^30^ (rs2532404) associated with progressive supranuclear palsy^31^ and was recently shown via CRISPRi to regulate *KANSL1* expression in iPSC-derived neurons^21^.

### Identification of AD-specific peak-gene-TF trios

To further investigate the regulatory roles of links, we identified peak-gene-TF trios in which: 1) there was a correlation between the linked peak and linked gene; 2) the accessibility of a linked peak harboring a specific TF motif was correlated with the expression of that TF; and 3) the expression of the TF was correlated with the expression of the linked gene (**Figure 4A**; *Methods*). This approach is similar to a recently described method called TRIPOD^32^. We performed these correlation analyses separately using either AD or control data sets to enable identification of TFs whose activities may be associated with disease. We restricted these analyses to links with a correlation value *>*0.3 that were within 100 kb of the linked gene’s TSS (115,107) and identified 60,120 peak-gene-TF trios involving 17,149 unique peaks and 437 TFs (**Table S6**). Fewer than 20% of the peaks in these trios are found in promoters, with the majority present in intronic regions (**Figure 4B**). Trio peaks were enriched for ENCODE distal (OR=1.26, p=2.2 × 10^−16^) and proximal (OR=1.12, p=5.9 × 10^−07^) enhancer-like sequences. There was a median of 37 trios per TF. The TF MEF2C was the most common trio participant, appearing in nearly 5% of all trios. While *MEF2C* was expressed in most cell types, expression of target genes in MEF2C trios were distinct between cell types (**Figures 4C and 4D**) In microglia, target genes were enriched in pattern recognition receptor (PRR) signaling and for synaptic transmission in neurons (**Figure 4E, Table S7**). PRRs consist of several receptor families including Toll-like receptors that are critical for microglial activation^33^.

**Figure 4.**
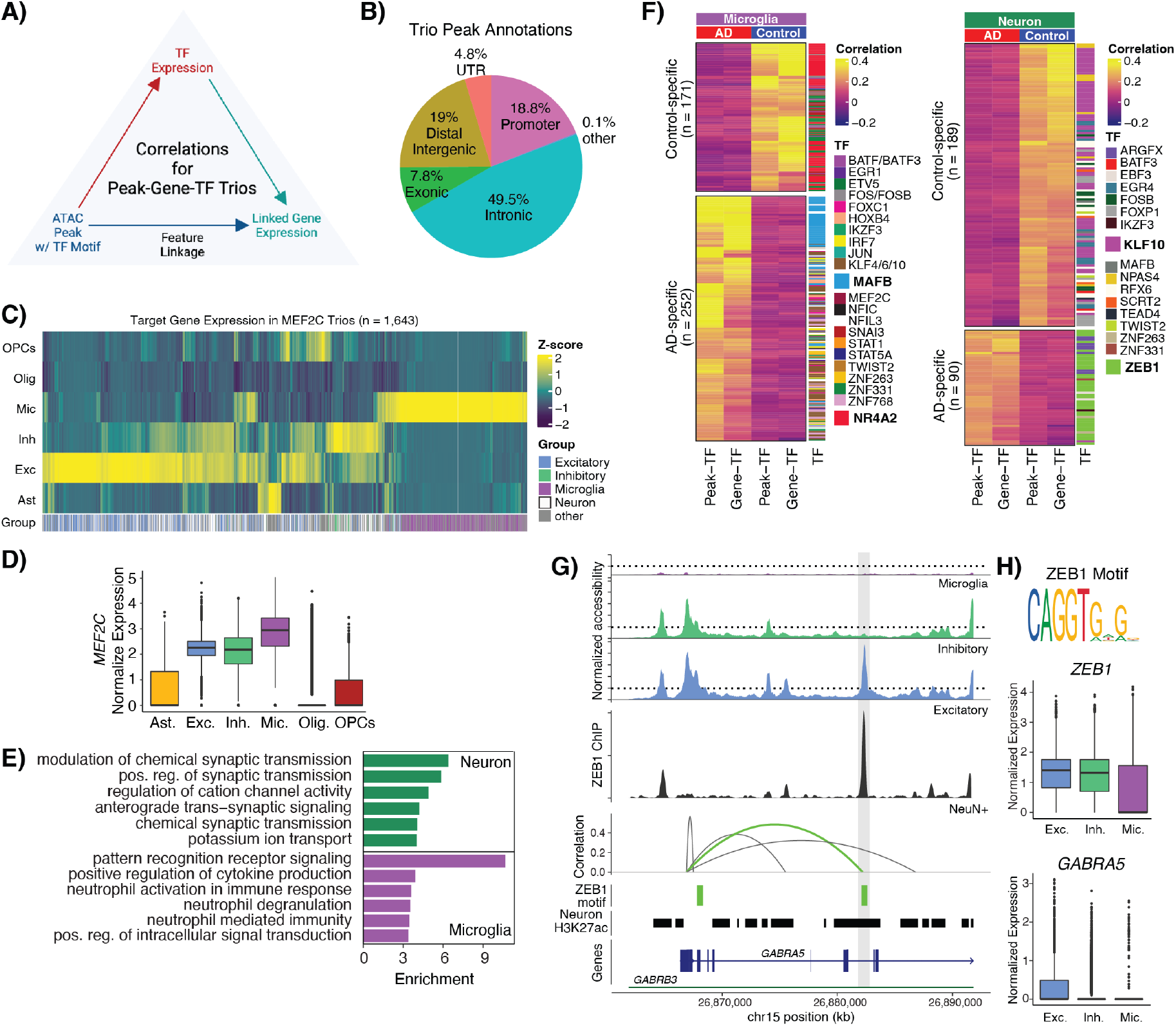
Identification of AD-specific TF regulatory networks. **A)** Strategy for defining peak-gene-TF trios. A linked peak containing a TF motif must be correlated with that TF and the expression of that TF must be correlated with the linked gene for that peak to be considered a trio. **B)** Genome annotations for location of linked peaks within trios. **C)** Heatmap of column-normalized expression of genes within MEF2C trios by cell type. **D)** Normalized expression of *MEF2C* by cell type. **E)** Top enriched gene ontology terms for genes within MEF2C trios from excitatory and inhibitory neurons (green = “Neuron”) and microglia (purple = “Microglia”). **F)** Heatmap of correlation values of AD and control-specific trios identified in microglia (left) and excitatory/inhibitory neurons (right). **G)** Linkage plot for *GABRA5*. Top panels: coverage plot of pseudo-bulked accessibility in indicated cell types. Middle panel: coverage plot of ZEB1 ChIP-seq signal from NeuN+ DLPFC tissue from two unaffected donors (1238 and 1242). Bottom panels: significant peak-gene links; green indicates overlap with ZEB1 motif. Arc height represents strength and direction of correlation. Track of ZEB1 motifs (green) and H3K27ac peaks from neurons (black; Nott et al^11^). Linked peak of interest is highlighted in gray. **H)** ZEB1 motif from JASPAR 2022 (top). Normalized expression of *ZEB1* and *GABRA5* in excitatory/inhibitory neurons and microglia.

Within this set of trios, there was a small subset that were specific to either AD or control groups (n = 2,718). While many of these were specific to a single cell type, 55% were shared across two or more (**Figure S4**). All cell type-specific trios overlapped H3K27ac peaks from their respective cell types (**Table S6**). Within microglia trios, NR4A2 was identified most frequently in the control-specific trios (**Figure 4F**). NR4A2 can function as both an activator and repressor and has been shown to repress inflammatory responses in microglia through recruitment of the CoREST complex^34,35^. Target genes in NR4A2 trios are enriched in neutrophil degranulation (OR = 9.01, q-value = 5.3 × 10^−6^) and include interleukin genes *IL1A* and *IL1B*, as well as *TGFB1*. Similarly, MAFB was involved in 24% of the AD-specific trios (**Figure 4F**) where it was linked to the microglial marker gene *CX3CR1* and genes involved in microglial activation (*TLR3, CD84, HAVCR2*)^36^. In healthy microglia, MAFB inhibits inflammatory responses^37^, consistent with our finding that target genes in AD-specific trios were enriched for negative regulation of myeloid leukocyte mediated immunity (OR = 332, q-value = 0.0004).

Within neuronal trios, we identified KLF10 and ZEB1 most frequently in control– and AD–specific trios, respectively (**Figure 4F**). These two TFs were also the most frequently observed in excitatory neuron trios; there were no inhibitory-specific trios identified (**Figure S4B**). In neurons, we identified ZEB1 in nearly 60% of all AD-specific trios with target genes involved in regulating ion channel signaling (*ITPR1, CAMK2A, CACNB3, KCNH3, KCNQ5*, and *KCNT1*). ZEB1 was never found in control-specific trios. Given the frequency of ZEB1 participation in neuronal AD-specific trios, we performed ZEB1 ChIP-seq in NeuN^+^ nuclei isolated from two control donors (1238 and 1242). We found that 41% of neuronal ZEB1 trios are bound by ZEB1, and these include 57 peaks within the AD-specific trios. The GABA_A_ receptor *α*5 subunit, encoded by *GABRA5*, is one gene that we find likely to be regulated by ZEB1 in AD (**Figure 4G**). *α*5 GABA_A_ receptors are associated with learning and memory, consistent with highest expression of *GABRA5* in hippocampal neurons and association of reduced expression with neurodevelopmental disorders^38^. In our data, *ZEB1* is expressed in both neurons and microglia; however, *GABRA5* is primarily expressed in excitatory neurons (**Figure 4G**, right). In excitatory neurons, we identified a linked peak correlated with *GABRA5* expression that was marked with H3K27ac and contained a ZEB1 motif. ChIP-seq data from two of our unaffected donors confirmed ZEB1 binding at this site providing additional evidence to suggest *cis* regulatory activity of this region for *GABRA5*.

### Genetic variation at candidate CREs

We performed stratified linkage disequilibrium score (sLDSC) regression^39^ to determine if our links were significantly enriched for SNPs associated with complex brain-related traits (**Figure 5A, Table S8**). Consistent with previous studies^40,41^, our microglia links were significantly enriched for heritability of AD across five different studies^2-6^; however, this was not true for those microglia links identified only in control samples, suggesting that variants in AD-specific CREs could have a greater contribution to AD risk. Specificity of microglia links for AD heritability is also supported by the lack of significant enrichment of these feature links with risk variants from other brain-related traits^42-46^ or traits where other immune cells play important roles^47-49^. In contrast, links identified in other cell types were enriched for heritability of brain-related traits including autism spectrum disorder (ASD), bipolar disorder (BD) and schizophrenia (SZ) with AD-specific links largely excluded from any significant enrichment in these traits. These findings are consistent with previous studies where candidate CREs identified in excitatory and inhibitory neurons were significantly associated with neuropsychiatric traits^11^. As expected, we identified no significant enrichments with immune diseases or with other phenotypic traits, such as body mass index (BMI)^50^ or height^51^.

**Figure 5.**
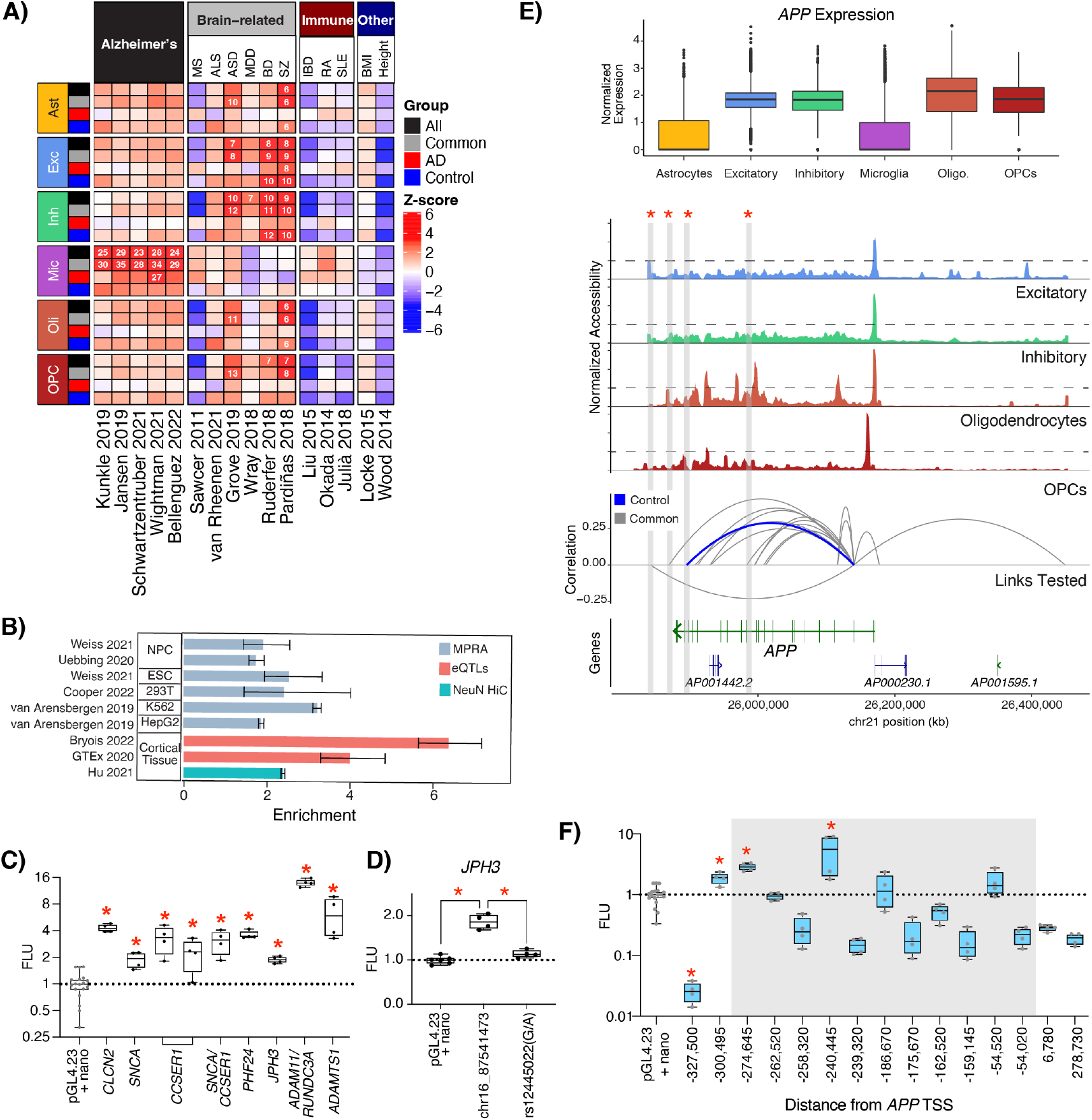
Identification of AD-specific TF regulatory networks. **A)** sLDSC results using 16 GWAS traits as indicated with our linked peaks stratified by cell type and group (“All” = all links, “Common” = links identified in both AD and control data, “AD” = links specific to AD, “Control” = links specific to control). Heatmap indicates coefficient z-score from running sLSDC with each set of links combined with the 97 baseline features. Feature-trait combinations with a z-score significantly larger than 0 (one-sided z-test with alpha = 0.05, P-values corrected within each trait using Holm’s method) are indicated with a numeric value reporting the enrichment score. **B)** Bar plot showing enrichment (+/- 95% CI) of feature linkages for previously nominated regulatory regions: active MPRA elements (blue), eQTLs where target gene is same as linked gene (pink), and HiC loops linking region to same target gene (green). **C)** Box plots showing statistically significant (* indicates p *<*0.05, ANOVA with Fisher’s LSD) elements representing feature linkages tested in luciferase assays. Luciferase elements are denoted by the linked gene for the nominated region. **D)** Box plots showing comparison of rs12445022 to its corresponding reference element linked to *JPH3* (* indicates p *<*0.05, ANOVA with Fisher’s LSD). **E)** Top: Normalized expression of *APP* in each cell type. Middle panels: Coverage plot of accessibility in indicated cell types. Bottom panel: Significant control (blue) and common (gray) gene-peak links to *APP* tested in luciferase assays. Arc height represents strength and direction of correlation. Links that contained CREs that increased expression of the luciferase reporter are highlighted in gray. F) Box plots showing all tested luciferase elements representing APP-peak links. Elements highlighted in gray are located within the *APP* gene body (* indicates p *<*0.05, ANOVA with Fisher’s LSD).

### Validation of candidate CREs

We compared the *>*300,000 links to pre-existing, large-scale functional genomic datasets to determine which candidate elements had previously shown evidence of regulatory activity. Three data types were considered to provide orthogonal evidence of regulatory activity: 1) massively parallel reporter assays (MPRAs)^21,52-54^, 2) eQTL studies^30,55^, and 3) HiC^56^ datasets. We found significant enrichments of links across each of these datasets despite several MPRAs being performed in cancer cell lines (**Figure 5B**). The MPRA data provided evidence that linked peaks could stimulate transcription, but are not capable of identifying the target gene. In contrast, HiC data from NeuN^+^ nuclei provided orthogonal validation of a linked peak’s target gene, but no evidence of promoting transcriptional activity. However, we intersected the results from these analyses and found that 1,542 of the 60,473 links that displayed regulatory activity in one or more MPRAs also identified the same target gene as the HiC data. In addition, 617 linked peaks overlapped eQTLs and were linked to the same gene providing both evidence of activity and confirming the target gene. Of the 67,541 links that overlapped at least one functional dataset, only 1,668 were also identified by Morabito et al.^19^.

For additional validation, we selected 51 neuronal links for testing in a luciferase reporter assay (**Table S9**). We performed these assays in the neuroepithelial-derived human embryonic kidney 293 (HEK293 and 293FT) cell lines because of the similar chromatin accessibility landscape to that found in brain tissues^21^. These cell lines are also technically tractable as they are highly transfectable and allow for efficient screening of regions of interest. We did not select any AD-specific links for validation, as we are using cell lines from a presumably unaffected individual. Thirteen of these 51 links contained SNPs associated with a brain-related trait (e.g. AD, epilepsy, neurodegeneration, etc.) and we tested both alleles of these SNPs (**Table S10**). Twelve of the elements increased activity of the luciferase reporter including regions linked to *SNCA* (*α*-synuclein) and *APP* (amyloid precursor protein) (**Figures 5C–5F**). Three of these active elements were involved in peak-gene-TF trios (*CCSER1*-MEF2C, *JPH3*-RARB, and *ADAMTS1*-SOX10). ChIP-seq analysis of NeuN^+^ nuclei confirmed that MEF2C is bound at the peak linked to *CCSER1*, a gene associated with autism^57^ (data not shown). Only one of the 15 variants tested abolished activity, rs12445022, a G/A substitution in a peak linked to *JPH3* (p = 0.0003 by ANOVA with Fisher’s LSD) (**Figure 5D**). *JPH3* encodes junctophilin-3, important for regulating neuronal excitability^58^. This *JPH3* linked peak was highly correlated (r = 0.64) with *JPH3* expression in both AD and control samples in all cell types except microglia. The linked peak is located 45,503 bp upstream of the *JPH3* TSS and was also linked to *ZCCHC14-DT*, although with a much lower correlation (r = 0.36). Repeat expansions in *JPH3* have been associated with a Huntington’s disease-like phenotype^59,60^.

Due to its importance in AD pathogenesis, we focus our validation efforts particulalry on the *APP* locus (**Figure 5E**) where we tested 15 elements and identified three that increased expression in the luciferase reporter assay (**Figure 5F**). *APP* is expressed across all cell types (**Figure 5E**, top panel) consistent with the high promoter accessibility observed (**Figure 5E**, middle panels). We also found one element with a negative correlation with *APP* expression that significantly reduced reporter activity; however, this assay was not designed to detect repressor activity and further experiments are required to assign a repressive function to this element.

## Discussion

Single cell multiomics has allowed for the generation of a rich source of disease– and cell type–specific candidate CREs enriched in variants associated with AD. Our study provides tangible advances by employing snRNA-seq and snATAC-seq in the same cells. Others have generated snRNA-seq and snATAC-seq separately and integrated them to identify CREs in AD^19^; however, profiling gene expression and chromatin accessibility simultaneously in the same nuclei allows for greater confidence in the correlations linking potential CREs to target genes. As such, we identified five times as many new candidate CREs (319,905 links vs 56,552 gene-linked cCREs) than previously reported^19^. To our knowledge, only one other study of another human neurodegenerative disease, Parkinson’s, used the 10X Genomics Multiomics (ATAC+Gene Expression) technology^61^ and identified a similarly large number of peak-gene linkages. Our approach is unique in that we identified peak-gene correlations independently in control and AD data sets allowing us to identify 40,831 peak-gene links specific to AD.

Our study provides two main advances in our understanding of altered gene regulation in AD. First, by leveraging the AD– and control–specific links identified here we constructed peak-gene-TF trios to determine which TFs were particularly involved in regulating AD-specific transcriptional programs. MAFB and ZEB1 were found to be enriched in AD-specific trios in microglia and neurons, respectively. MAFB has been implicated in exercise-associated responses in the peripheral immune system in AD^62^. Centrally, MAFB has been implicated in regulation of the receptor VISTA in microglia, which is up-regulated in AD^63^. In this study, we identify a previously unknown role for ZEB1 in AD-specific transcriptional regulation. Previously, ZEB1 was shown to play a critical role in epithelial-mesenchymal transition in neural crest migration and glioblastoma^64,65^ and further investigation is necessary to reveal its role in AD. Secondly, we demonstrated enhancer-like activity for 12 candidate CREs linked to neurodegeneration-associated genes *APP, SNCA, PHF24, ADAM11*, and *ADAMTS1*. This study lays the groundwork for additional functional validation in future studies to confirm these genes as targets of these CREs.

One limitation of this study is that snATAC-seq data can contain spurious signals, as well as bias from transcribed genes. This limitation underscores the importance of evaluation via orthogonal methods, which we have provided using both published and newly generated data. A second limitation is that our sample size is small. This can be addressed in future studies by increasing sample size; however, the shared signals we observed with larger AD snRNA-seq studies emphasizes the representative nature of our sample set, and that our total number of cells per biological sample is adequate. Finally, as with any study from postmortem tissue, we are measuring by definition the material that remains in a neurodegenerative disease, which can confound interpretation. For this reason, we chose to evaluate DLPFC, which is preserved later into the disease course of AD than tissues affected earlier such as entorhinal cortex and hippocampus.

In summary, our study provides important new insights into the contribution of CREs to AD including the roles of TFs ZEB1 and MAFB in neurons and microglia. These findings could provide additional insights for interpreting SNPs associated with AD risk should they disrupt binding motifs for these TFs. Further, these TFs could be therapeutic targets for manipulating aberrant gene regulation in AD. Our study lays the groundwork for future research to expand on the candidate– and literature-based validation approaches taken here. High throughput CRISPRi screens are well-suited to test the necessity and sufficiency of regulatory elements for linked gene expression. Future validation efforts will greatly contribute to advancing our understanding of the effects of non-coding variation on risk for AD.

## Supporting information

Description of Supplemental Tables

## Author contributions

Conceptualization, L.F.R, J.N.C, and J.M.L; Formal Analysis, A.G.A., L.F.R., and I.R.; Investigation, J.N.B, L.M.W., S.C.R., L.F.R., J.M.L., and B.B.R.; Data Curation, A.G.A.; Writing –Original Draft, L.F.R., A.G.A., and B.B.R.; Writing –Review and Editing, J.N.C., R.M.M., and J.M.L.; Supervision, L.F.R., J.N.C., and R.M.M; Funding, J.N.C. and R.M.M.

## Declaration of interests

The authors declare no competing interests.

## Methods

### Resource availability

Further information and requests for resources and reagents should be directed to and will be fulfilled by the lead contact Lindsay Rizzardi.

### Experimental model and subject details

#### Cell cultures

HEK293 cells were obtained from ATCC (CRL-1573) and grown in DMEM (high glucose, L-glutamine, no sodium pyruvate) (ThermoFisher), supplemented with 10% fetal bovine serum (FBS). 293FT cells were obtained from ThermoFisher Scientific (R70007) and maintained in DMEM (high glucose, L-Glutamine, 100 mg/L Sodium Pyruvate) supplemented with 10% FBS, 1% Glutamax, 1% non-essential amino acids (NEAA), and 500 mg/mL Geneticin (G418 Sulfate, ThermoFisher). All cells were cultured at 37^*◦*^C with 5% CO2.

#### Human brain tissues

Postmortem human brain biospecimens were obtained from the NIH Neurobiobank at the University of Miami and the Human Brain and Spinal Fluid Resource Center (HBSFRC) and from collaborators from the Pritzker Neuropsychiatric Disorders Research Consortium in the Department of Psychiatry and Human Behavior, University of California Irvine (UCI) as noted in **Table S1**. Flash-frozen tissues were obtained from the dorsolateral prefrontal cortex (BA9/46) of 9 donors diagnosed with Alzheimer’s (Braak stages 4-6) and 9 unaffected controls. Demographic information for each donor is presented in Table S1. No statistical methods were used to pre-determine sample sizes, but our sample sizes are similar to those reported in previous publications^18,19^. Data collection and analyses were not performed blind to tissue of origin. We did not pre-select samples based on *APOE* genotype, but genotype information was generated for each sample through TaqMan genotyping assays (see *APOE Genotyping*).

### Method details

#### Nuclei Isolation from human brain tissues for single nucleus multiomics

Approximately 50-100 mg of frozen tissue per sample was homogenized in 4 mL of nuclei extraction buffer [0.32 M sucrose, 10 mM Tris - pH 7.4, 5 mM CaCl_2_, 3 mM Mg(Ac)_2_, 1 mM DTT, 0.1 mM EDTA, 0.1% Triton X-100, 0.2U/*µ*L Protector RNAse inhibitor (Sigma cat. 3335399001)] by douncing 30 times in a 40 mL dounce homogenizer. Filter through 70 *µ*m filter and spin at 500x*g*, 5 min at 4^*◦*^C in a swinging bucket centrifuge. Resuspend nuclei in 500 *µ*L nuclei extraction buffer and layer over 750 *µ*L sucrose solution (1.8 M sucrose, 10 mM Tris pH 7.4, 3 mM Mg(Ac)_2_, 1 mM DTT) in a 1.5 mL tube. The samples were then centrifuged at *>*16,000x*g* for 30 min at 4^*◦*^C. After centrifugation, the supernatant was removed by aspiration and the nuclear pellet was resuspended in 125 *µ*L PBS with 1% BSA and centrifuged 5 min at 500 x g at 4^*◦*^C in a swinging bucket centrifuge. Permeabilization was performed according to 10X Genomics protocol CG000375 Rev B: nuclei were resuspended in 100 *µ*L lysis buffer (10 mM Tris-pH 7.4,10 mM NaCl, 3 mM MgCl_2_, 1% BSA, 0.01% Tween-20, 0.01% NP-40, 0.001% digitonin, 1 mM DTT, 1 U/*µ*L Protector RNase inhibitor) and incubated 2 min on ice. Nuclei were washed once and resuspended in 30 *µ*L of 1X nuclei buffer with 1 mM DTT and 0.5 U/*µ*L of Protector RNAse inhibitor. Nuclei quality and concentrations were determined using the Countess II FL (ThermoFisher).

#### Single nucleus multiomics

Transposition, nuclei isolation, barcoding, and library preparation were performed according to the 10X Genomics Chromium Next GEM Single Cell Multiome protocol CG000338 Rev E with the following alterations. The initial set of eight samples were processed as above (noted as “batch 1” in **Table S1**) and each sample was loaded across two lanes of the Chromium Next GEM Chip J. Nuclei were loaded according to manufacturer’s recommendations to target recovery of 10,000 nuclei per lane. The second batch of ten samples were processed as above, but two samples were pooled per lane of the Chromium Next GEM Chip J (each pool is indicated by sub-batch in **Table S1**). Each pool consisted of a male and female donor to facilitate assignment of each single cell back to the donor based on genotype and chrY gene expression (see *Sample Demultiplexing*). For these samples, we pooled 20,000 nuclei from each sample and the entire pool was processed according to the multiome protocol. Libraries were sequenced by HudsonAlpha Discovery using Illumina NovaSeq S4 flowcells.

#### Sample demultiplexing

For lanes where a male and female sample were pooled together, reads were assigned to samples by genotyping cells. Variants were called from the cellranger output bam file for each cell using cellsnp-lite^66^. High-confidence SNPs from the 1000 Genome Project were used as a reference panel to call variants. Cell genotypes were then split by individual using vireoSNP with the number of donors set to two ^67^. Cells were labeled as donor_0, donor_1, unassigned, or doublet. Unassigned and doublet cells were removed. Donor ID was assigned to the sample by observing the number of UMIs for genes on chrY. The donor ID with the higher mean counts was assigned to the male sample (**Table S11**).

#### Joint snRNA-seq and snATAC-seq workflow

Low-quality cells were filtered on gene expression data (nFeatures *>* 200, nFeatures *<* 10,000, and mitochondrial percent *<* 5) and chromatin accessibility data (nucleosome signal *<* 2 and TSS enrichment *>* 2). PMI-associated genes^68^ were removed from the RNA counts matrix. Peaks that were present in less than 10 cells were removed from the ATAC matrix. RNA counts were normalized with SCTransform with mitochondrial percent per cell regressed out. Principal component analysis (PCA) was performed on RNA, and UMAP was run on the first 30 principal components (PCs). The optimum number of PCs was determined to be 30 PCs using an elbow plot. The ATAC counts were normalized with term-frequency inverse-document-frequency (TFIDF).

Dimension reduction was performed with singular value decomposition (SVD) of the normalized ATAC matrix. The ATAC UMAP was created using the 2^nd^ through the 50^th^ LSI components. Doublet density was computed using computeDoubletDensity from scDblFinder where doublet score is the ratio of densities of simulated doublets to the density in the data. Cells with a doublet score *>* were removed. Normalization and dimension reduction were performed again on the filtered set with the same parameters. Predicted cell types were determined for each cell using Seurat SCT-normalized reference mapping. Gene expression data was mapped to SCT-normalized DLPFC data^12^ and annotated with the cell types of the reference map. Cells with a predicted cell type score *<* 0.95 were removed from the data. Batch effects were corrected in RNA (theta=1) and ATAC (theta=2) with Harmony (v1.0.0)^69^ by removing the effect of sample.

#### WNN analysis of snRNA-seq and snATAC-seq

The weighted nearest neighbor (wnn) graph was determined with Seurat’s *FindMultiModalNeighbors* to represent a weighted combination of both modalities. The first 30 dimensions of the Harmony-corrected RNA reduction and the 2^nd^ through the 50^th^ dimensions from the Harmony-corrected ATAC reduction were used to create the graph. The WNN UMAP was created using the wknn (k=20) (**Figure S5**).

#### Differential expression

Differentially expressed genes (DEGs) were determined for AD versus control for each cell type. Within each cell type, the gene expression data was log-normalized with a scale factor of 1 × 10^5^. Pericytes and Endothelial cells were not included in the analysis because of small cell counts.

Differential expression was assessed using MAST^70^ for genes present in at least 25% of either AD or control cells. Age and sex were included as covariates in the MAST model. Genes with a Bonferroni-adjusted p-value *<* 0.01 and an absolute log2 fold change *>* 0.25 were determined to be significant. DEGs between cell types were determined using MAST with age and sex as covariates for genes present in at least 25% of cells. Genes with a Bonferroni adjusted p-value *<* 0.01 and an absolute log2 fold change *>* 0.5 were determined to be significant.

#### Annotation of cell subpopulations

Cell type subclusters were identified using weighted snRNA and snATAC modalities. Expression data were normalized with SCTransform, and chromatin accessibility data were normalized with TFIDF within each cell type. Normalized values were used to construct a multimodal weighted nearest neighbor graph (k=20). Clusters were identified using wknn and the SLM algorithm. The resolution (0.3, 0.2, 0.3, 0.3, 0.45) was adjusted for each cell type (Astro, Inh, Exc, Olig, Mic). Any cluster with *<* 100 cells was excluded from DEG analysis. Within each cell type, cluster DEGs were determined for each subcluster versus all other subclusters. DEGs were defined as those with a Bonferroni adjusted p-value *<*0.01 using MAST with age and sex as covariates. Only genes that were detected in at least 25% of cells in a subcluster were considered.

Neuronal subclusters were further annotated with Azimuth ^22^ Human motor cortex ^71^ clusters to identify known neuronal subpopulations. For each neuronal subcluster, a subtype was assigned by the enrichment for up-regulated subcluster DEGs in Azimuth gene sets. Enrichment was performed using enrichR^72-74^ and the Azimuth Cell Types 2021 gene sets. The top subtype annotation was assigned to a subcluster if the adjusted p-value was *<* 0.01.

AD-specific subclusters and subtypes were determined by observing overrepresentation of cells isolated from AD individuals. Statistically significant overrepresentation was evaluated with a Fisher’s exact test and adjusted p-values.

#### Gene set enrichment

The R package enrichR^72-74^ was used for all gene set enrichment analyses. Sets of DEGs and peak-linked genes were used as input to look for enrichment in GO Biological Process 2021, GO Molecular Function 2021, GO Cellular Component 2021, and KEGG 2021 databases. Terms with an adjusted p-value *<* 0.05 were considered to be enriched.

#### Feature linkage analysis

ATAC peaks were called independently for each cell type using MACS2 and Signac *CallPeaks* and the union of these peaks was used in subsequent analyses retaining the cell type annotations. The peaks were then annotated with ChIPseeker^75^ and *TxDb*.*Hsapiens*.*UCSC*.*hg38*.*knownGene* where promoters were considered to be 1 kb upstream and 100 bp downstream of the TSS. Only ATAC peaks that were present in at least 2% of cells in at least one cell type were included in linkage analyses. AD and control linkages were identified separately via the cellranger-arc (v2.0) reanalyze function using the filtered cell type ATAC peaks and either AD or control expression and accessibility data as input. The maximum interaction distance was restricted to 500 kb. Peak-peak links are produced by the cellranger-arc pipeline by default, but were not used for downstream analyses. Feature linkages with an absolute correlation score *>* 0.2 and linked to a gene with *<* 200 UMIs were removed.

#### Gene-Peak-TF trios

Trios were called for a filtered set of feature linkages by removing links further than 100 kb and links with an absolute score *<* 0.2. Motifs were then called in each linked-peak using Signac *AddMotifs* and the JASPAR 2022 ^76^ CORE PFM. Peaks with *>* 100 motifs were additionally filtered from the link set. TF expression, linked-gene expression, and linked-peak accessibility matrices for trio correlation were derived from the average counts within metacells. Metacells were determined using WNN clusters for all AD cells and all control cells separately. TF-peak scores are the Pearson correlation between peak accessibility and the expression of the TF whose motif was called in the peak. TF-gene scores are the Pearson correlation between a gene and the TF whose motif was called in the linked-peak. Significant associations were defined as those with a p-value *<* 0.001. Significant trios were then defined as those with a significant positive TF-peak correlation and a significant TF-gene correlation.

#### Partitioned heritability analysis

To evaluate whether feature links are enriched with common genetic variants that have been associated with AD or other traits by GWAS, we performed stratified linkage disequilibrium (LD) score regression (sLDSC v1.0.1)^39,77^. sLDSC estimates the proportion of genome-wide SNP-based heritability that can be attributed to SNPs within a given genomic feature by a regression model that combines GWAS summary statistics with estimates of LD from an ancestry-matched reference panel. Summary statistics for AD were downloaded from^2–6^. To estimate SNP heritability from AD GWAS summary statistics, we excluded the *APOE* and *MHC/HLA* genomic regions. Additional GWAS summary statistics were downloaded for brain-related^42-46,78^ and other traits^47-51^. Each cell type feature link was tested individually along with the full baseline model (baseline-LD model v2.2.) that included 97 categories capturing a broad set of genomic annotations. Links to GWAS summary statistics are available in **Table S8**. Additional files needed for the sLDSC analysis were downloaded from https://alkesgroup.broadinstitute.org/LDSCORE/ following instructions at https://github.com/bulik/ldsc/wiki.

#### *APOE* genotyping

To determine *APOE* status, TaqMan genotyping assays (cat#: 4371353) were used to genotype SNPs rs429358 and rs7412 (cat: 4351379, C___3084793_20 and C___904973_10, respectively) following the manufacturer’s instructions. Genotyping calls were made using QuantStudio software (v1.3) for all individuals in this study. *APOE* status is reported in **Table S1**.

#### Comparisons to external data sources

Cell type-specific H3K27ac peak calls were obtained from^11^ and converted to hg38 coordinates using the *liftOver* function from the R package rtracklayer. GABA and GLU neuronal sub-type H3K27ac fastqs from Kozlenkov et al. ^28^ were downloaded from Synapse (syn12033252) and processed as individual replicates using the AQUAS Transcription Factor and Histone ChIP-Seq processing pipeline^79^. (https://github.com/kundajelab/chipseq_pipeline). Peaks were called using the IDR naive overlapping method with a threshold of 0.05 and the optimal peak sets were used. For each cell type, only peaks identified in at least 3 individuals were retained for downstream analyses. ATAC-seq peaks from non-neuronal cell types were intersected with H3K27ac data from the corresponding cell type obtained from Nott et al.^11^. Excitatory and inhibitory neuron ATAC-seq peaks were intersected with H3K27ac peaks identified from GLU (NeuN^+^/SOX6^-^) or GABA (NeuN^+^/SOX6^+^) neuronal nuclei^28^ and from neuronal (NeuN^+^) nuclei^11^. MPRA data was obtained from^21,52-54^, eQTL data was obtained from^30,55^, and neuronal HiC loop calls were obtained from^56^.

#### Plasmids

The pNL1.1.CMV [Nluc/CMV] and pGL4.23 [luc2/minP] vectors were obtained from Promega. Luciferase elements were generated by selecting 467 bp of the nominated region using hg38 coordinates. Both the forward and reverse complement sequences were ordered as gBlocks from Integrated DNA Technologies (IDT). Gibson assembly was performed by cloning elements into the pGL4.23 [luc2/minP] vector digested with *EcoRV*. Element insertion was confirmed by Sanger sequencing (MCLAB). Each element was individually prepped 3 times for a total of 6 individual plasmid preparations per nominated region.

#### Transfection

HEK293 and 293FT cells were plated at 70,000 cells/cm^2^ in a 24-well format. Before plating 293FT cells, culture plates were pre-coated with poly-L-ornithine solution (Millipore Sigma). The next day, cells were transfected with 1 *µ*g of plasmid DNA using Lipofectamine LTX with Plus Reagent (ThermoFisher) following the manufacturer’s recommendations. Per transfection, 900 ng of luciferase element and 100 ng of pNL1.1.CMV [Nluc/CMV] were used. A transfection reaction of 900 ng pGL4.23 [luc2/minP] and 100 ng pNL1.1.CMV [Nluc/CMV] was used as a baseline control. Both vectors were also transfected as background controls (100 ng) with pmaxGFP (900 ng, Lonza). Cell lysates were harvested by freezing at -80^*◦*^C 48 hours post-transfection.

#### Luciferase assays

Luciferase assays were performed using the Nano-Glo Dual-Luciferase Reporter Assay System (Promega) following the manufacturer’s protocol. Cell lysis was performed on the 24 well plate and aliquoted across 4 wells of a white 96-well plate for 4 technical replicates per biological replicate.

Assays were completed in quadruplicate. Firefly luminescence was first normalized across the average plate luminescence and then normalized to the average control luminescence. For each biological replicate, the median fold luminescence value was determined for the four technical replicates. Four biological replicates were compared to the pGL4.23 [luc2/minP]/pNL1.1.CMV [Nluc/CMV] control using an ordinary one-way ANOVA with Fisher’s LSD.

#### Chromatin preparation for sorted nuclei

Buffers required: Nuclei Extraction Buffer (NEB): 0.32 M Sucrose, 5 mM CaCl_2_, 3 mM Mg(Ac)_2_, 0.1 mM EDTA, 10 mM Tris-HCl, 0.1 mM PMSF, 0.1% Triton X-100, 1 mM DTT. Before use, add Roche cOmplete protease inhibitor cocktail according to manufacturer recommendation (Sigma 11697498001). Sucrose Cushion Buffer (SCB): 1.6 M Sucrose, 3 mM Mg(Ac)_2_, 10 mM Tris-HCl, 1 mM DTT. Interphase Buffer: 0.8 M Sucrose, 3 mM Mg(Ac)_2_, 10 mM Tris-HCl. Blocking buffer: 1x PBS, 1% BSA, 1 mM EDTA. Pellet buffer: add up to 200 *µ*L 1 M CaCl_2_ to 10 mL SCB. RIPA: 1x PBS, 1% NP-40, 0.5% sodium deoxycholate, 0.1% SDS.

Methods for extracting and sorting nuclei from postmortem brain are similar to previously published methods^80^. Here, approximately 500 mg of tissue was placed into a chilled 40 mL Dounce homogenizer containing 5 mL of NEB on ice and allowed to partially thaw to ease douncing (2-3 minutes). Extract nuclei by douncing with “tight” pestle 30-40 times until the tissue is homogenized. Transfer to 15 mL conical tube on ice, wash glassware with 5 mL NEB and add to 15 mL tube. Fix chromatin by adding 625 *µ*L of 16% formaldehyde (methanol free, Thermo 28906) to a final concentration of 1% and rotate end-over-end at room temperature for 10 minutes. Halt fixation by adding 500 *µ*L of 2.5 M Glycine and incubate another 5 minutes rotating at room temperature then place homogenate back on ice. During fixation, prepare sucrose gradient in two ultracentrifuge buckets (Beckman Coulter cat:344058) by layering 5 mL of Interphase buffer atop 10 mL of SCB in each. Carefully layer nuclei homogenate atop sucrose gradient, balance with NEB, then ultracentrifuge at 24,000 rpm for 2 h using SW28 swinging bucket rotor (Beckman Coulter). Upon completion, inspect tubes for a visible pellet of nuclei at the bottom of tube. Remove debris at interphase first by using a 25 mL graduated pipette, then continue removing the remaining sucrose gradient being careful not to disturb the nuclei pellet. Carefully resuspend the pellet in 1 mL cold PBS and transfer to a 15 mL lo-bind tube containing 2 mL PBS on ice. (Optional: if pellet appears to contain large debris then pass through 70 *µ*m filter). Wash ultracentrifuge tubes with 1 mL cold PBS and combine in 15 mL tube to a final volume of 10 mL, inverting to mix. Centrifuge the nuclei at 1,000xg for 10 minutes at 4^*◦*^C to remove residual sucrose. Label nuclei by resuspending pellet in 5 mL blocking buffer with NeuN-488 antibody (Millipore, cat: MAB377X) and OLIG2 antibody (Abcam, cat: ab109186) at 1:5,000 each. Incubate nuclei in staining buffer with rotation for at least 1 hour at 4^*◦*^C. Spin nuclei 500x*g* for 5 minutes to pellet, remove supernatant, then resuspend in 5 mL blocking buffer with goat-anti-rabbit-647 (Thermo, cat: A-21245) at 1:5,000 and DAPI at 1:100,000. Incubate for at least 1 hour at 4^*◦*^C with rotation. Remove stain by centrifuging 500x*g* 5 minutes at 4^*◦*^C and resuspending in 3 mL cold PBS. Hold on ice and proceed immediately to sorting.

Nuclei were sorted using Sony MA900 with a 70 *µ*m nozzle and pressure not exceeding pressure setting of 7. Gates were set to capture those populations that were positive for 488 signal (NeuN^+^), positive for 647 signal (OLIG^+^), or negative for both (NeuN^-^;OLIG^-^). Each population was collected into 5 mL tubes held at 4^*◦*^C and pooled into 15 mL lo-bind tubes on ice. Purity of selected samples were typically *>*95% based on reanalysis of sorted samples. To concentrate nuclei for downstream analysis, add approximately 2 mL of pellet buffer per 10 mL of sorted nuclei and rotated at 4^*◦*^C for 15 minutes. Centrifuge 500x^g^ for 10 minutes at 4^*◦*^C, after which a pellet should be visible. Remove supernatant and carefully resuspend pelleted nuclei in at least 3 mL cold PBS. Centrifuge 500x*g* for 5 minutes at 4^*◦*^C.

To generate chromatin for ChIP-seq, resuspend pellet in cold RIPA plus protease inhibitor (Roche, Sigma 11836153001) at approximately 3 million nuclei per 250 *µ*L. Transfer 250 *µ*L of each sample to the Bioruptor (Diagenode, cat: C30010016) tubes and sonicate tissue using a Bioruptor Pico (8 cycles; 30 seconds on/ 30 seconds off). Pool the sonicated chromatin into a 1.5 mL DNA lo-bind conical tube and centrifuge 12,000x*g* for 5 minutes at 4^*◦*^C to remove any insoluble debris. Collect supernatant into a separate tube, add RIPA to final volume equivalent to 500,000 nuclei per 100 *µ*L, then dispense working aliquots into 1.5mL tubes held on dry ice. Store at -80^*◦*^C.

#### ChIP-seq protocol

ChIP-seq for ZEB1 was performed using chromatin from NeuN^+^ sorted DLPFC nuclei from two control donors serving as biological replicates. ChIP-seq for MEF2C was performed on bulk DLPFC tissues from two control donors serving as biological replicates. Protocols for ChIP-seq are similar to those for frozen tissue previously described by our lab ^81,82^ and consistent with techniques recommended by the ENCODE Consortium (www.encodeproject.org/documents). Briefly, 60 *µ*L Dynabeads (ThermoFisher, cat: 11203D) were washed with cold 1x PBS + 5 mg/mL BSA then combined with 8 *µ*L antibody targeting ZEB1 (Bethyl, cat: A301-921A) or MEF2C (proteintech, cat: 18290-1-AP) in a final volume of 200 *µ*L and held at 4^*◦*^C overnight with rotation. Tubes of aliquoted chromatin are thawed on ice and bead/antibody complex is washed with PBS + 5 mg/mL BSA solution. Beads are ultimately resuspended in 100 *µ*L RIPA and bro*µ*ght to 200 *µ*L with 100 *µ*L chromatin aliquot. Incubate bead/antibody with chromatin using rotation for one hour at room temperature then move to 4^*◦*^C for another hour. After incubation, bead complexes were washed five times with a LiCl wash buffer (100 mM Tris at pH 7.5, 500 mM LiCl, 1% NP-40, 1% sodium deoxycholate) and wash with 1 mL of cold TE (10 mM Tris-HCl at pH 7.5, 0.1 mM Na_2_EDTA).

Chromatin was eluted from beads by incubating with intermittent shaking for 1 hour at 65^*◦*^C in IP elution buffer (1% SDS, 0.1 M NaHCO_3_), followed by incubating overnight at 65^*◦*^C to reverse formaldehyde cross-links. DNA was purified using DNeasy Blood and Tissue kit (Qiagen 69506) and eluted in a final volume of 50 *µ*L EB. Recovered DNA was quantified using Qubit dsDNA HS Assay kit (Thermo Q32854). For input controls, one aliquot of each tissue was brought to 200 *µ*L with RIPA and reverse-crosslinked overnight at 65^*◦*^C. The following morning, samples were incubated an additional 30 minutes with 20 *µ*L Proteinase K and 4 *µ*L RNase A (Qiagen 19101) and subsequently eluted for DNA using DNeasy Blood and Tissue kit. The entirety of the remaining IP DNA (and approximately 250 ng Input control) were used to generate sequencing libraries. Libraries were prepared by blunting and ligating ChIP DNA fragments to sequencing adapters for amplification with barcoded primers (30 sec at 98^*◦*^C; [10 sec at 98^*◦*^C, 30 sec at 65^*◦*^C, 30 sec at 72^*◦*^C] x 15 cycles; 5 min at 72^*◦*^C). Libraries were quantified with Qubit dsDNA HS Assay kit and visualized with Standard Sensitivity NGS Fragment Analysis Kit (Advanced Analytical DNF-473) and Fragment Analyzer 5200 (Agilent). Libraries were sequenced using Illumina NovaSeq flowcell with 100 bp single-end runs.

#### ChIP-seq analysis

Prior to analysis, reads were processed to remove optical duplicates with clumpify (BBMap v38.20; https://sourceforge.net/projects/bbmap/) [dedupe=t optical=t dupedist=2500] and remove adapter reads with Cutadapt (v1.16)^83^ [-a AGATCGGAAGAGC -m 40]. Input reads were capped at 40 million using Seqtk (v1.2; https://github.com/lh3/seqtk). Individual experiments were constructed following ENCODE guidelines (https://www.encodeproject.org/about/experiment-guidelines/) and analyzed with the chip-seq-pipeline2 processing pipeline (https://github.com/ENCODE-DCC/chip-seq-pipeline2). All software within the package was run using the default settings or those recommended by the authors for transcription factors. Final peaks were called using the IDR naïve overlapping method with a threshold of 0.05.

## Data and code availability

The raw and processed data generated will be made available through NCBI GEO under series accession number GSE214637 upon publication. All supplementary tables are available upon request. All the code generated during this study is available at aanderson54/scMultiomics_AD

## Acknowledgments

We thank Paige I. Hall for assisting L.W. and J.N.B., and Rebecca M. Hauser for *APOE* genotyping. This study was supported in part by NIH grant 3R01MH110472-03S1 awarded to R.M.M. and NIH grant 5R00AG068271 awarded to J.N.C. Additional funding was generously provided by donors to the HudsonAlpha Foundation Memory and Mobility Fund. We especially thank the donors and their families without whom this work would not be possible.

## Supporting Information

**Figure S1.**
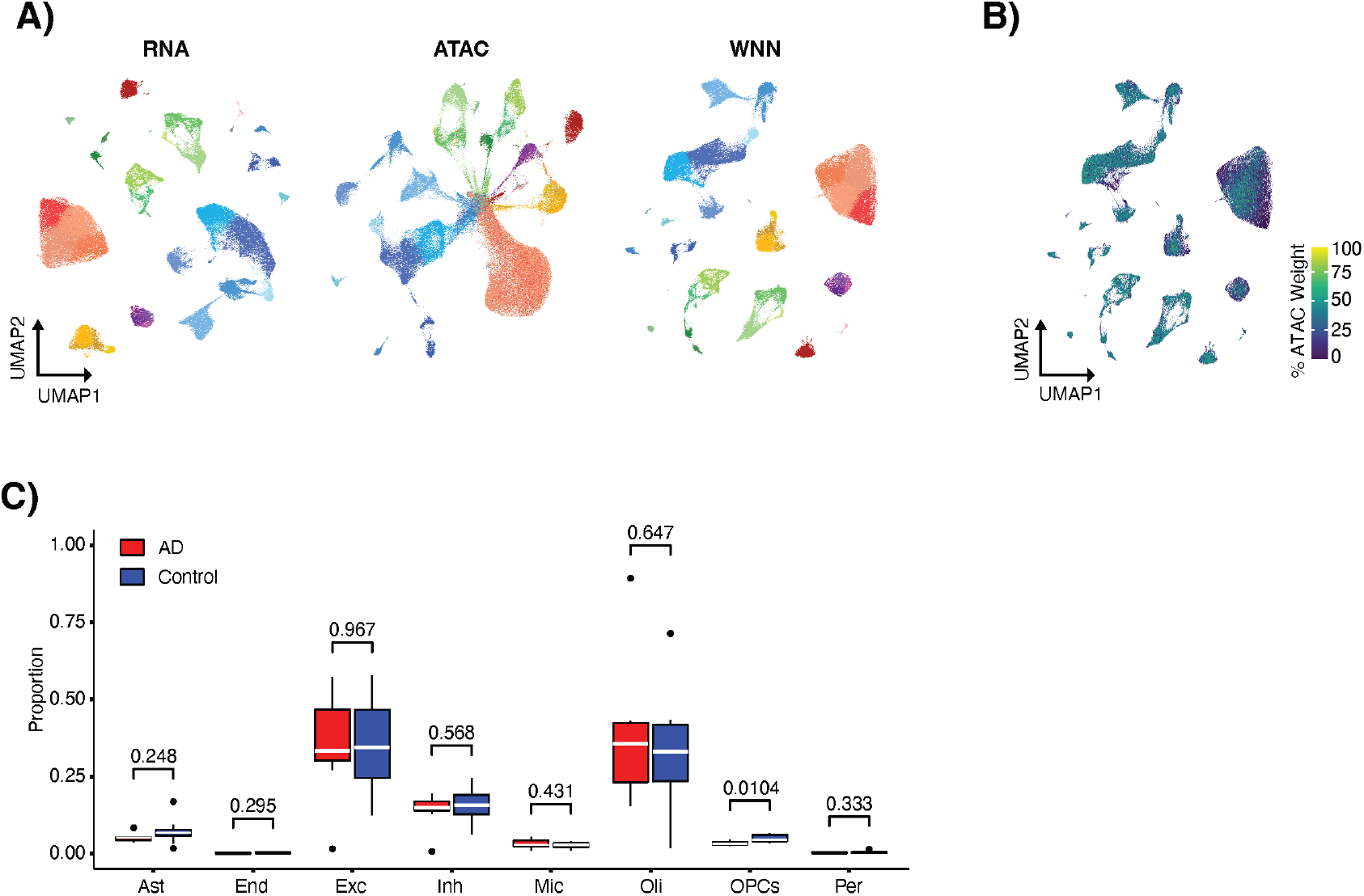
Integrating snRNA-seq and snATAC-seq data. **A)** UMAP visualization of cells represented by only snRNA-seq data, only snATAC-seq data, and joint WNN. Cells are colored by cell type and cluster assignment. **B)** WNN UMAP colored by the percent weight given to the snATAC-seq data for each cell when creating the WNN graph. **C)** The proportion of cells assigned to a cell type from each individual. P-values from t-test are indicated above box plots.

**Figure S2.**
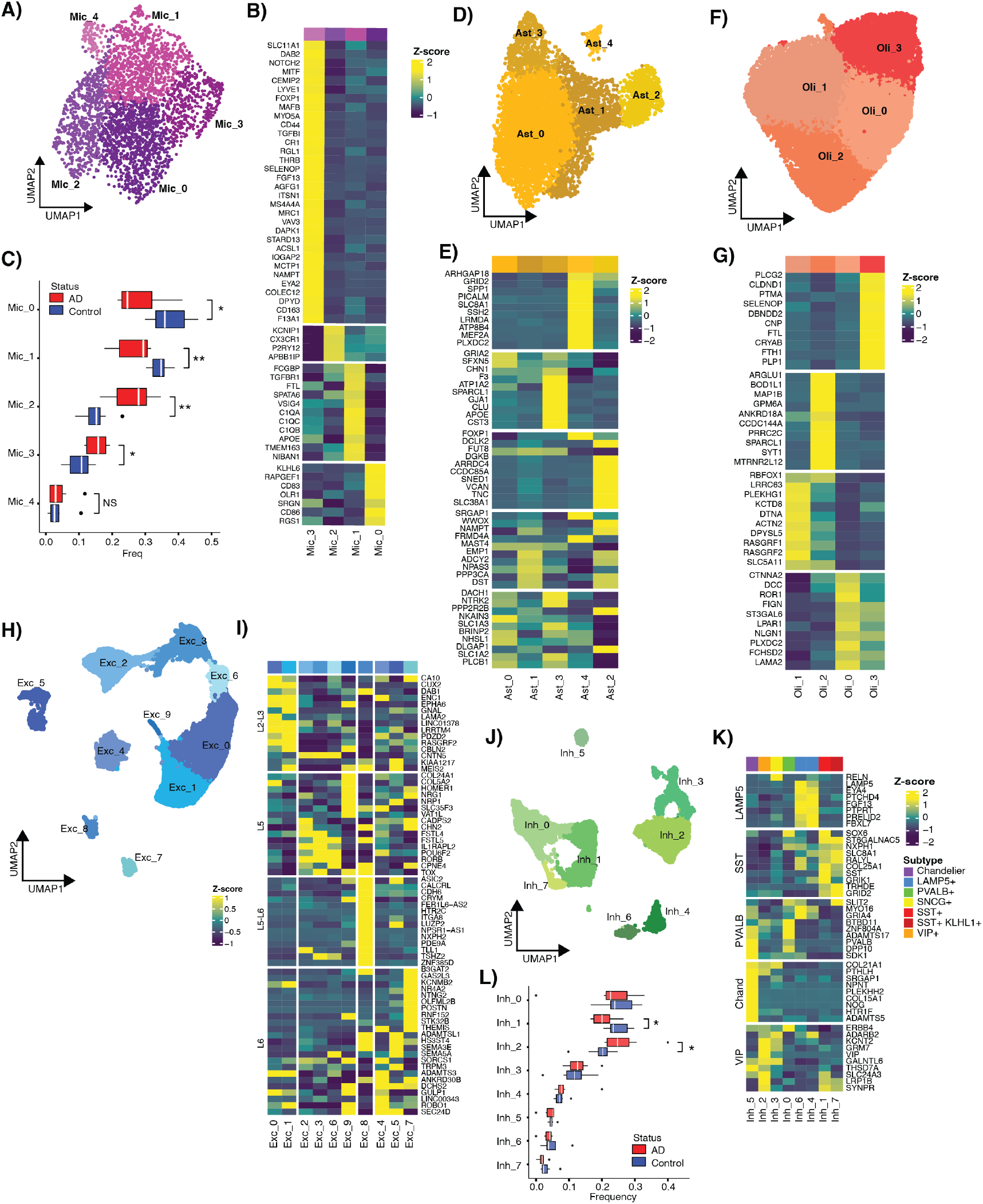
Identification of cell type subclusters. **A)** UMAP visualization of 5 microglia subclusters.**B)** Heatmap of row-normalized expression for the top DEGs for each microglia subcluster. **C)** The proportion of cells assigned to each subcluster from each individual (* indicates subclusters with a t-test p-value *<* 0.05; ** p-value *<* 0.01). **D)** UMAP visualization of 5 astrocyte subclusters. **E)** Heatmap of row-normalized expression for the top 10 DEGs for each astrocyte subcluster. **F)** UMAP visualization of the 4 oligodendrocyte subclusters. **G)** Heatmap of row-normalized expression for the top 10 DEGs for each oligodendrocyte subcluster. **H)** UMAP visualization of the 10 excitatory neuron subclusters. **I)** Heatmap of row-normalized expression for Azimuth Glutamatergic subtype markers. **J)** UMAP visualization of the 8 inhibitory subclusters. **K)** Heatmap of row-normalized expression for Azimuth GABAergic subtype markers. **L)** The proportion of cells assigned to each inhibitory subcluster from each individual (* indicated t-test p-value *<* 0.05).

**Figure S3.**
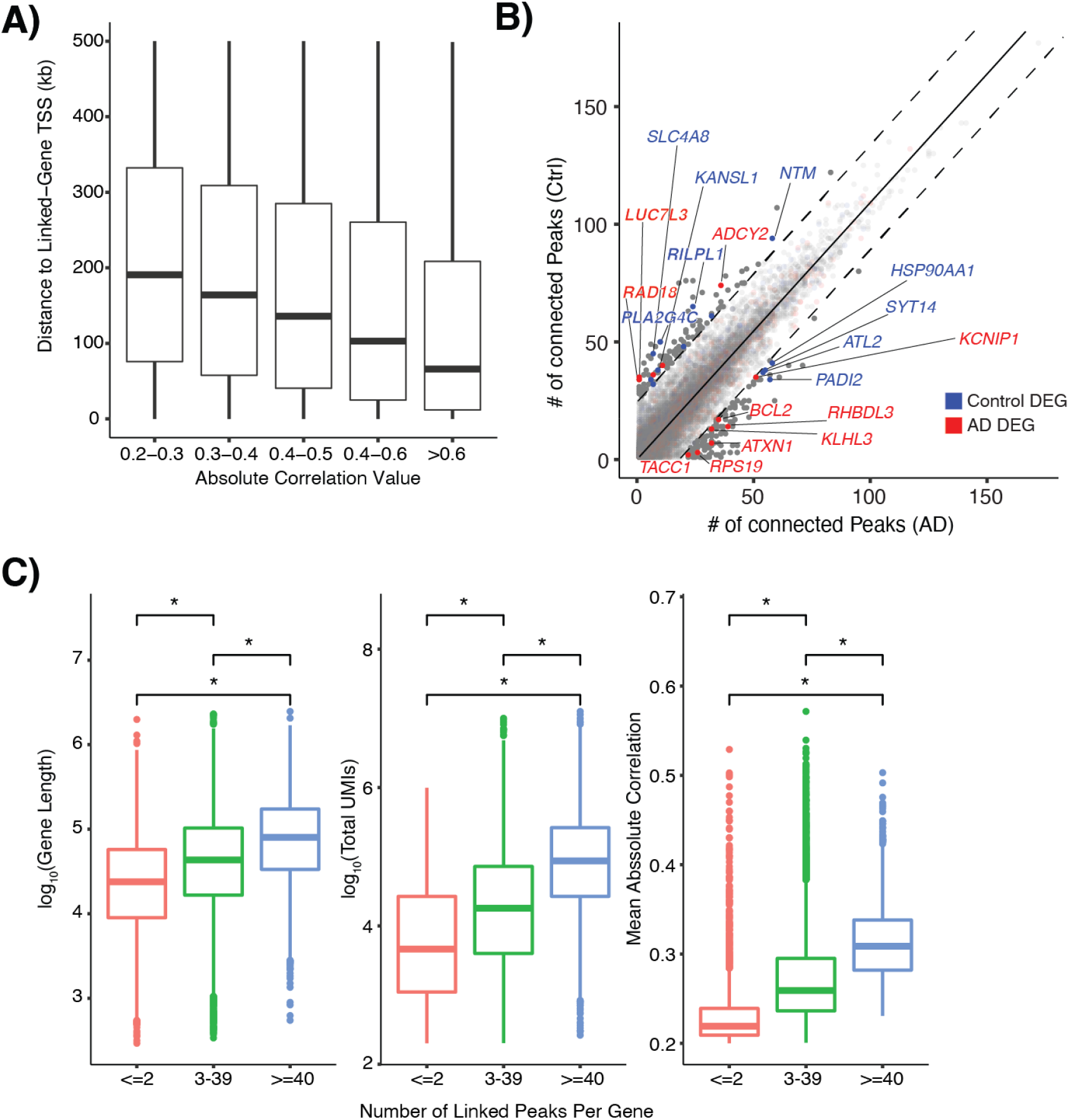
Feature linkage description. **A)** Characteristics of genes by number of links. Left panel: distribution of gene length by the number of links to the gene. Middle panel: distribution of UMIs by the number of links to the gene. Right panel: average absolute correlation score by the number of links to the gene. **B)** Distribution of the distance to linked-gene TSS by binned absolute correlation.

**Figure S4.**
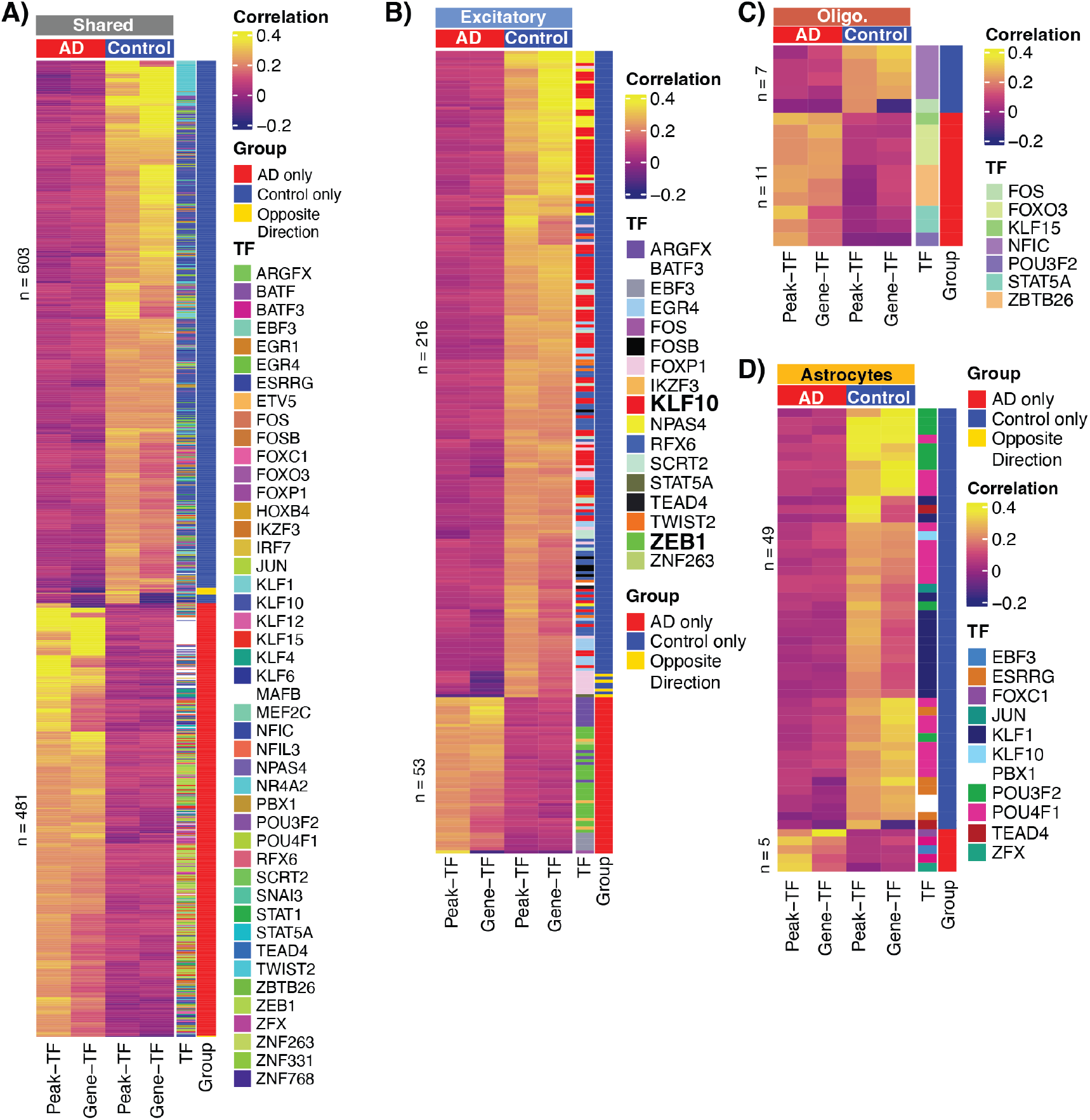
AD and control-specific peak-gene-TF trios. **A)** Heatmap of correlation values of AD and control specific trios identified in links shared across cell types, **B)** excitatory neurons, **C)** oligodendrocytes, and **D)** astrocytes.

**Figure S5.**
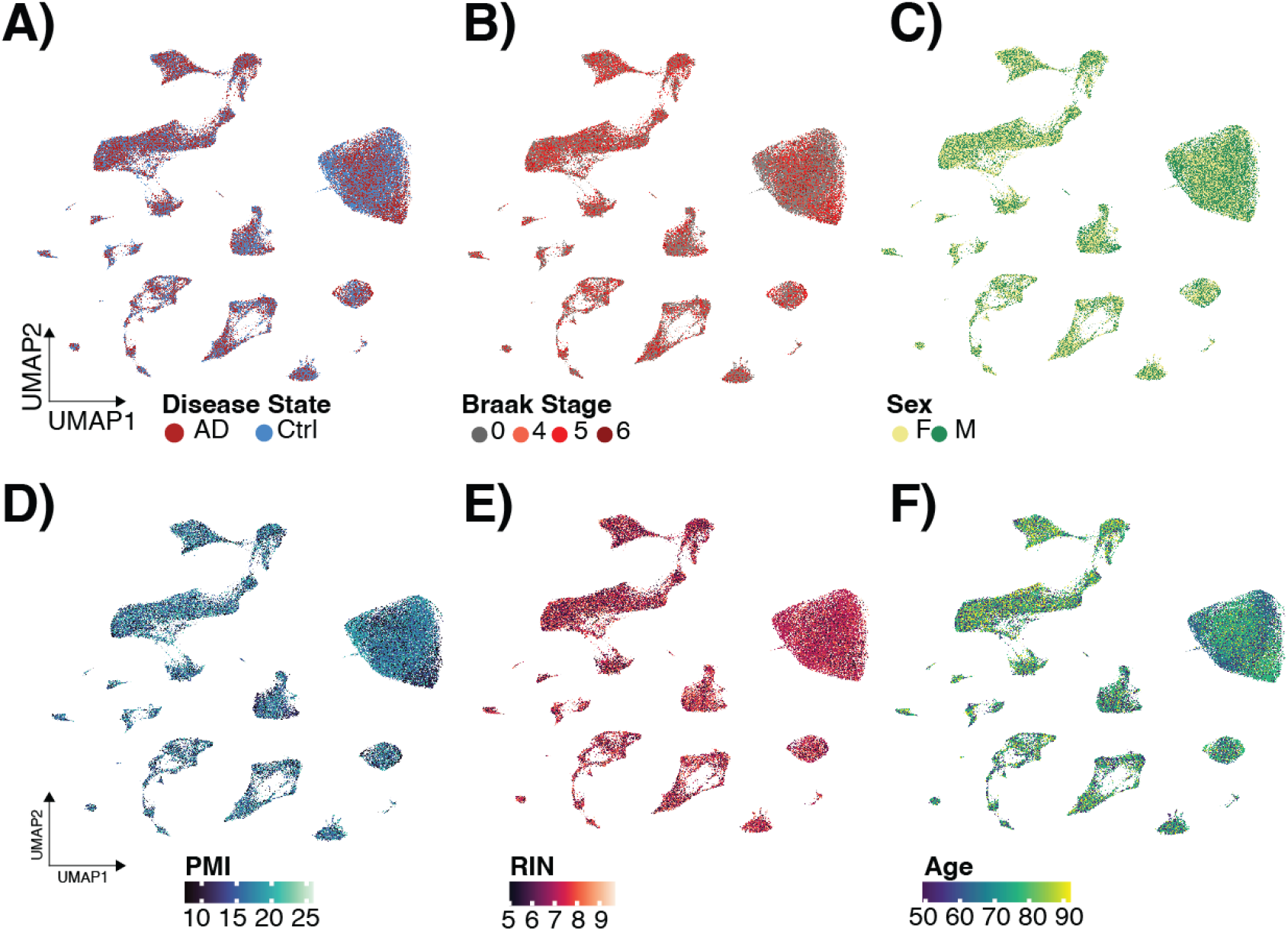
Donor characteristics across cell types. WNN UMAP colored by **A)** disease status, **B)** Braak Stage, **C)** sex, **D)** postmortem interval (PMI), **E)** RNA integrity number (RIN), and **F)** Age.

## Notes

### Competing Interest Statement

The authors have declared no competing interest.

## References

1. Neuner, S.M., Tcw, J., and Goate, A.M. (2020). Genetic architecture of Alzheimer’s disease. Neurobiol. Dis. 143, 104976. 10.1016/j.nbd.2020.104976.

2. Kunkle, B.W., Grenier-Boley, B., Sims, R., Bis, J.C., Damotte, V., Naj, A.C., Boland, A., Vronskaya, M., van der Lee, S.J., Amlie-Wolf, A., et al. (2019). Genetic meta-analysis of diagnosed Alzheimer’s disease identifies new risk loci and implicates Aβ, tau, immunity and lipid processing. Nat. Genet. 51, 414–430. 10.1038/s41588-019-0358-2.

3. Jansen, I.E., Savage, J.E., Watanabe, K., Bryois, J., Williams, D.M., Steinberg, S., Sealock, J., Karlsson, I.K., Hägg, S., Athanasiu, L., et al. (2019). Genome-wide meta-analysis identifies new loci and functional pathways influencing Alzheimer’s disease risk. Nat. Genet. 9, 63. 10.1038/s41588-018-0311-9.

4. Schwartzentruber, J., Cooper, S., Liu, J.Z., Barrio-Hernandez, I., Bello, E., Kumasaka, N., Young, A.M.H., Franklin, R.J.M., Johnson, T., Estrada, K., et al. (2021). Genome-wide meta-analysis, fine-mapping and integrative prioritization implicate new Alzheimer’s disease risk genes. Nat. Genet., 1–11. 10.1038/s41588-020-00776-w.

5. Wightman, D.P., Jansen, I.E., Savage, J.E., Shadrin, A.A., Bahrami, S., Holland, D., Rongve, A., Børte, S., Winsvold, B.S., Drange, O.K., et al. (2021). A genome-wide association study with 1,126,563 individuals identifies new risk loci for Alzheimer’s disease. Nat. Genet. 53, 1276–1282. 10.1038/s41588-021-00921-z.

6. Bellenguez, C., Küçükali, F., Jansen, I.E., Kleineidam, L., Moreno-Grau, S., Amin, N., Naj, A.C., Campos-Martin, R., Grenier-Boley, B., Andrade, V., et al. (2022). New insights into the genetic etiology of Alzheimer’s disease and related dementias. Nat. Genet. 54, 412–436. 10.1038/s41588-022-01024-z.

7. Uffelmann, E., Huang, Q.Q., Munung, N.S., de Vries, J., Okada, Y., Martin, A.R., Martin, H.C., Lappalainen, T., and Posthuma, D. (2021). Genome-wide association studies. Nat. Rev. Methods Primer 1, 1–21. 10.1038/s43586-021-00056-9.

8. Zhang, F., and Lupski, J.R. (2015). Non-coding genetic variants in human disease. Hum. Mol. Genet. 24, R102–R110. 10.1093/hmg/ddv259.

9. Khani, M., Gibbons, E., Bras, J., and Guerreiro, R. (2022). Challenge accepted: uncovering the role of rare genetic variants in Alzheimer’s disease. Mol. Neurodegener. 17, 3. 10.1186/s13024-021-00505-9.

10. Corces, M.R., Shcherbina, A., Kundu, S., Gloudemans, M.J., Frésard, L., Granja, J.M., Louie, B.H., Eulalio, T., Shams, S., Bagdatli, S.T., et al. (2020). Single-cell epigenomic analyses implicate candidate causal variants at inherited risk loci for Alzheimer’s and Parkinson’s diseases. Nat. Genet. 52, 1158–1168. 10.1038/s41588-020-00721-x.

11. Nott, A., Holtman, I.R., Coufal, N.G., Schlachetzki, J.C.M., Yu, M., Hu, R., Han, C.Z., Pena, M., Xiao, J., Wu, Y., et al. (2019). Brain cell type-specific enhancer-promoter interactome maps and disease risk association. Science 366, 1134–1139. 10.1126/science.aay0793.

12. Mathys, H., Davila-Velderrain, J., Peng, Z., Gao, F., Mohammadi, S., Young, J.Z., Menon, M., He, L., Abdurrob, F., Jiang, X., et al. (2019). Single-cell transcriptomic analysis of Alzheimer’s disease. Nature 570, 332–337. 10.1038/s41586-019-1195-2.

13. Grubman, A., Chew, G., Ouyang, J.F., Sun, G., Choo, X.Y., McLean, C., Simmons, R.K., Buckberry, S., Vargas-Landin, D.B., Poppe, D., et al. (2019). A single-cell atlas of entorhinal cortex from individuals with Alzheimer’s disease reveals cell-type-specific gene expression regulation. Nat. Neurosci. 22, 2087–2097. 10.1038/s41593-019-0539-4.

14. Sadick, J.S., O’Dea, M.R., Hasel, P., Dykstra, T., Faustin, A., and Liddelow, S.A. (2022). Astrocytes and oligodendrocytes undergo subtype-specific transcriptional changes in Alzheimer’s disease. Neuron 110, 1788-1805.e10. 10.1016/j.neuron.2022.03.008.

15. Leng, K., Li, E., Eser, R., Piergies, A., Sit, R., Tan, M., Neff, N., Li, S.H., Rodriguez, R.D., Suemoto, C.K., et al. (2021). Molecular characterization of selectively vulnerable neurons in Alzheimer’s disease. Nat. Neurosci. 24, 276–287. 10.1038/s41593-020-00764-7.

16. Zhou, Y., Song, W.M., Andhey, P.S., Swain, A., Levy, T., Miller, K.R., Poliani, P.L., Cominelli, M., Grover, S., Gilfillan, S., et al. (2020). Human and mouse single-nucleus transcriptomics reveal TREM2-dependent and TREM2-independent cellular responses in Alzheimer’s disease. Nat. Med. 26, 131–142. 10.1038/s41591-019-0695-9.

17. Alsema, A.M., Jiang, Q., Kracht, L., Gerrits, E., Dubbelaar, M.L., Miedema, A., Brouwer, N., Hol, E.M., Middeldorp, J., van Dijk, R., et al. (2020). Profiling Microglia From Alzheimer’s Disease Donors and Non-demented Elderly in Acute Human Postmortem Cortical Tissue. Front. Mol. Neurosci. 13.

18. Lau, S.-F., Cao, H., Fu, A.K.Y., and Ip, N.Y. (2020). Single-nucleus transcriptome analysis reveals dysregulation of angiogenic endothelial cells and neuroprotective glia in Alzheimer’s disease. Proc. Natl. Acad. Sci. 117, 25800–25809. 10.1073/pnas.2008762117.

19. Morabito, S., Miyoshi, E., Michael, N., Shahin, S., Martini, A.C., Head, E., Silva, J., Leavy, K., Perez-Rosendahl, M., and Swarup, V. (2021). Single-nucleus chromatin accessibility and transcriptomic characterization of Alzheimer’s disease. Nat. Genet. 53, 1143–1155. 10.1038/s41588-021-00894-z.

20. Kartha, V.K., Duarte, F.M., Hu, Y., Ma, S., Chew, J.G., Lareau, C.A., Earl, A., Burkett, Z.D., Kohlway, A.S., Lebofsky, R., et al. (2022). Functional inference of gene regulation using single-cell multi-omics. Cell Genomics 2. 10.1016/j.xgen.2022.100166.

21. Cooper, Y.A., Teyssier, N., Dräger, N.M., Guo, Q., Davis, J.E., Sattler, S.M., Yang, Z., Patel, A., Wu, S., Kosuri, S., et al. (2022). Functional regulatory variants implicate distinct transcriptional networks in dementia. Science 377, eabi8654. 10.1126/science.abi8654.

22. Hao, Y., Hao, S., Andersen-Nissen, E., Mauck, W.M., Zheng, S., Butler, A., Lee, M.J., Wilk, A.J., Darby, C., Zager, M., et al. (2021). Integrated analysis of multimodal single-cell data. Cell 184, 3573-3587.e29. 10.1016/j.cell.2021.04.048.

23. Stuart, T., Srivastava, A., Madad, S., Lareau, C.A., and Satija, R. (2021). Single-cell chromatin state analysis with Signac. Nat. Methods 18, 1333–1341. 10.1038/s41592-021-01282-5.

24. Belonwu, S.A., Li, Y., Bunis, D., Rao, A.A., Solsberg, C.W., Tang, A., Fragiadakis, G.K., Dubal, D.B., Oskotsky, T., and Sirota, M. (2022). Sex-Stratified Single-Cell RNA-Seq Analysis Identifies Sex-Specific and Cell Type-Specific Transcriptional Responses in Alzheimer’s Disease Across Two Brain Regions. Mol. Neurobiol. 59, 276–293. 10.1007/s12035-021-02591-8.

25. DeTomaso, D., and Yosef, N. (2021). Hotspot identifies informative gene modules across modalities of single-cell genomics. Cell Syst. 12, 446-456.e9. 10.1016/j.cels.2021.04.005.

26. Fulco, C.P., Nasser, J., Jones, T.R., Munson, G., Bergman, D.T., Subramanian, V., Grossman, S.R., Anyoha, R., Doughty, B.R., Patwardhan, T.A., et al. (2019). Activity-by-contact model of enhancer–promoter regulation from thousands of CRISPR perturbations. Nat. Genet. 51, 1664–1669. 10.1038/s41588-019-0538-0.

27. ENCODE Project Consortium, Moore, J.E., Purcaro, M.J., Pratt, H.E., Epstein, C.B., Shoresh, N., Adrian, J., Kawli, T., Davis, C.A., Dobin, A., et al. (2020). Expanded encyclopaedias of DNA elements in the human and mouse genomes. Nature 583, 699–710. 10.1038/s41586-020-2493-4.

28. Kozlenkov, A., Li, J., Apontes, P., Hurd, Y.L., Byne, W.M., Koonin, E.V., Wegner, M., Mukamel, E.A., and Dracheva, S. (2018). A unique role for DNA (hydroxy)methylation in epigenetic regulation of human inhibitory neurons. Sci. Adv. 4, eaau6190. 10.1126/sciadv.aau6190.

29. Zollino, M., Orteschi, D., Murdolo, M., Lattante, S., Battaglia, D., Stefanini, C., Mercuri, E., Chiurazzi, P., Neri, G., and Marangi, G. (2012). Mutations in KANSL1 cause the 17q21.31 microdeletion syndrome phenotype. Nat. Genet. 44, 636–638. 10.1038/ng.2257.

30. Bryois, J., Calini, D., Macnair, W., Foo, L., Urich, E., Ortmann, W., Iglesias, V.A., Selvaraj, S., Nutma, E., Marzin, M., et al. (2022). Cell-type-specific cis-eQTLs in eight human brain cell types identify novel risk genes for psychiatric and neurological disorders. Nat. Neurosci. 25, 1104–1112. 10.1038/s41593-022-01128-z.

31. Chen, J.A., Chen, Z., Won, H., Huang, A.Y., Lowe, J.K., Wojta, K., Yokoyama, J.S., Bensimon, G., Leigh, P.N., Payan, C., et al. (2018). Joint genome-wide association study of progressive supranuclear palsy identifies novel susceptibility loci and genetic correlation to neurodegenerative diseases. Mol. Neurodegener. 13, 41. 10.1186/s13024-018-0270-8.

32. Jiang, Y., Harigaya, Y., Zhang, Z., Zhang, H., Zang, C., and Zhang, N.R. (2022). Nonparametric Interrogation of Transcriptional Regulation in Single-Cell RNA and Chromatin Accessibility Multiomic Data. 2021.09.22.461437. 10.1101/2021.09.22.461437.

33. Kigerl, K.A., de Rivero Vaccari, J.P., Dietrich, W.D., Popovich, P.G., and Keane, R.W. (2014). Pattern recognition receptors and central nervous system repair. Exp. Neurol. 258, 5–16. 10.1016/j.expneurol.2014.01.001.

34. Saijo, K., Winner, B., Carson, C.T., Collier, J.G., Boyer, L., Rosenfeld, M.G., Gage, F.H., and Glass, C.K. (2009). A Nurr1/CoREST transrepression pathway attenuates neurotoxic inflammation in activated microglia and astrocytes. Cell 137, 47. 10.1016/j.cell.2009.01.038.

35. K, S., B, W., Ct, C., Jg, C., L, B., Mg, R., Fh, G., and Ck, G. (2009). A Nurr1/CoREST pathway in microglia and astrocytes protects dopaminergic neurons from inflammation-induced death. Cell 137. 10.1016/j.cell.2009.01.038.

36. Patir, A., Shih, B., McColl, B.W., and Freeman, T.C. (2019). A core transcriptional signature of human microglia: Derivation and utility in describing region-dependent alterations associated with Alzheimer’s disease. Glia 67, 1240–1253. 10.1002/glia.23572.

37. Holtman, I.R., Skola, D., and Glass, C.K. (2017). Transcriptional control of microglia phenotypes in health and disease. J. Clin. Invest. 127, 3220–3229. 10.1172/JCI90604.

38. Jacob, T.C. (2019). Neurobiology and Therapeutic Potential of α5-GABA Type A Receptors. Front. Mol. Neurosci. 12.

39. Finucane, H.K., Bulik-Sullivan, B., Gusev, A., Trynka, G., Reshef, Y., Loh, P.R., Anttila, V., Xu, H., Zang, C., Farh, K., et al. (2015). Partitioning heritability by functional annotation using genome-wide association summary statistics. Nat Genet 47, 1228–1235. 10.1038/ng.3404.

40. Tansey, K.E., Cameron, D., and Hill, M.J. (2018). Genetic risk for Alzheimer’s disease is concentrated in specific macrophage and microglial transcriptional networks. Genome Med. 10, 14. 10.1186/s13073-018-0523-8.

41. Novikova, G., Kapoor, M., Tcw, J., Abud, E.M., Efthymiou, A.G., Chen, S.X., Cheng, H., Fullard, J.F., Bendl, J., Liu, Y., et al. (2021). Integration of Alzheimer’s disease genetics and myeloid genomics identifies disease risk regulatory elements and genes. Nat. Commun. 12, 1610. 10.1038/s41467-021-21823-y.

42. International Multiple Sclerosis Genetics Consortium, Wellcome Trust Case Control Consortium 2, Sawcer, S., Hellenthal, G., Pirinen, M., Spencer, C.C.A., Patsopoulos, N.A., Moutsianas, L., Dilthey, A., Su, Z., et al. (2011). Genetic risk and a primary role for cell-mediated immune mechanisms in multiple sclerosis. Nature 476, 214–219. 10.1038/nature10251.

43. van Rheenen, W., van der Spek, R.A.A., Bakker, M.K., van Vugt, J.J.F.A., Hop, P.J., Zwamborn, R.A.J., de Klein, N., Westra, H.-J., Bakker, O.B., Deelen, P., et al. (2021). Common and rare variant association analyses in amyotrophic lateral sclerosis identify 15 risk loci with distinct genetic architectures and neuron-specific biology. Nat. Genet. 53, 1636–1648. 10.1038/s41588-021-00973-1.

44. Grove, J., Ripke, S., Als, T.D., Mattheisen, M., Walters, R.K., Won, H., Pallesen, J., Agerbo, E., Andreassen, O.A., Anney, R., et al. (2019). Identification of common genetic risk variants for autism spectrum disorder. Nat. Genet. 51, 431–444. 10.1038/s41588-019-0344-8.

45. Wray, N.R., Ripke, S., Mattheisen, M., Trzaskowski, M., Byrne, E.M., Abdellaoui, A., Adams, M.J., Agerbo, E., Air, T.M., Andlauer, T.M.F., et al. (2018). Genome-wide association analyses identify 44 risk variants and refine the genetic architecture of major depression. Nat. Genet. 50, 668–681. 10.1038/s41588-018-0090-3.

46. Pardiñas, A.F., Holmans, P., Pocklington, A.J., Escott-Price, V., Ripke, S., Carrera, N., Legge, S.E., Bishop, S., Cameron, D., Hamshere, M.L., et al. (2018). Common schizophrenia alleles are enriched in mutation-intolerant genes and in regions under strong background selection. Nat. Genet. 50, 381–389. 10.1038/s41588-018-0059-2.

47. Liu, J.Z., van Sommeren, S., Huang, H., Ng, S.C., Alberts, R., Takahashi, A., Ripke, S., Lee, J.C., Jostins, L., Shah, T., et al. (2015). Association analyses identify 38 susceptibility loci for inflammatory bowel disease and highlight shared genetic risk across populations. Nat. Genet. 47, 979–986. 10.1038/ng.3359.

48. Okada, Y., Wu, D., Trynka, G., Raj, T., Terao, C., Ikari, K., Kochi, Y., Ohmura, K., Suzuki, A., Yoshida, S., et al. (2014). Genetics of rheumatoid arthritis contributes to biology and drug discovery. Nature 506, 376–381. 10.1038/nature12873.

49. Julià, A., López-Longo, F.J., Pérez Venegas, J.J., Bonàs-Guarch, S., Olivé, À., Andreu, J.L., Aguirre-Zamorano, M. À., Vela, P., Nolla, J.M., de la Fuente, J.L.M., et al. (2018). Genome-wide association study meta-analysis identifies five new loci for systemic lupus erythematosus. Arthritis Res. Ther. 20, 100. 10.1186/s13075-018-1604-1.

50. Locke, A.E., Kahali, B., Berndt, S.I., Justice, A.E., Pers, T.H., Day, F.R., Powell, C., Vedantam, S., Buchkovich, M.L., Yang, J., et al. (2015). Genetic studies of body mass index yield new insights for obesity biology. Nature 518, 197–206. 10.1038/nature14177.

51. Wood, A.R., Esko, T., Yang, J., Vedantam, S., Pers, T.H., Gustafsson, S., Chu, A.Y., Estrada, K., Luan, J., Kutalik, Z., et al. (2014). Defining the role of common variation in the genomic and biological architecture of adult human height. Nat. Genet. 46, 1173–1186. 10.1038/ng.3097.

52. Weiss, C.V., Harshman, L., Inoue, F., Fraser, H.B., Petrov, D.A., Ahituv, N., and Gokhman, D. (2021). The cis-regulatory effects of modern human-specific variants. eLife 10, e63713. 10.7554/eLife.63713.

53. Uebbing, S., Gockley, J., Reilly, S.K., Kocher, A.A., Geller, E., Gandotra, N., Scharfe, C., Cotney, J., and Noonan, J.P. (2021). Massively parallel discovery of human-specific substitutions that alter enhancer activity. Proc. Natl. Acad. Sci. 118. 10.1073/pnas.2007049118.

54. van Arensbergen, J., Pagie, L., FitzPatrick, V.D., de Haas, M., Baltissen, M.P., Comoglio, F., van der Weide, R.H., Teunissen, H., Võsa, U., Franke, L., et al. (2019). High-throughput identification of human SNPs affecting regulatory element activity. Nat. Genet. 51, 1160–1169. 10.1038/s41588-019-0455-2.

55. Consortium, T. Gte. (2020). The GTEx Consortium atlas of genetic regulatory effects across human tissues. Science 369, 1318–1330. 10.1126/science.aaz1776.

56. Hu, B., Won, H., Mah, W., Park, R.B., Kassim, B., Spiess, K., Kozlenkov, A., Crowley, C.A., Pochareddy, S., Li, Y., et al. (2021). Neuronal and glial 3D chromatin architecture informs the cellular etiology of brain disorders. Nat. Commun. 12, 3968. 10.1038/s41467-021-24243-0.

57. Ruzzo, E.K., Pérez-Cano, L., Jung, J.-Y., Wang, L., Kashef-Haghighi, D., Hartl, C., Singh, C., Xu, J., Hoekstra, J.N., Leventhal, O., et al. (2019). Inherited and de novo genetic risk for autism impacts shared networks. Cell 178, 850-866.e26. 10.1016/j.cell.2019.07.015.

58. Garbino, A., van Oort, R.J., Dixit, S.S., Landstrom, A.P., Ackerman, M.J., and Wehrens, X.H.T. (2009). Molecular evolution of the junctophilin gene family. Physiol. Genomics 37, 175–186. 10.1152/physiolgenomics.00017.2009.

59. Bourinaris, T., Athanasiou, A., Efthymiou, S., Wiethoff, S., Salpietro, V., and Houlden, H. (2021). Allelic and phenotypic heterogeneity in Junctophillin-3 related neurodevelopmental and movement disorders. Eur. J. Hum. Genet. 29, 1027–1031. 10.1038/s41431-021-00866-1.

60. Schneider, S.A., Marshall, K.E., Xiao, J., and LeDoux, M.S. (2012). JPH3 Repeat Expansions Cause a Progressive Akinetic-Rigid Syndrome with Severe Dementia and Putaminal Rim in a Five-Generation African-American Family. Neurogenetics 13, 133–140. 10.1007/s10048-012-0318-9.

61. Adams, L., Song, M.K., Tanaka, Y., and Kim, Y.-S. (2022). Single-nuclei paired multiomic analysis of young, aged, and Parkinson’s disease human midbrain reveals age- and disease-associated glial changes and their contribution to Parkinson’s disease. 2022.01.18.22269350. 10.1101/2022.01.18.22269350.

62. Chen, Y., Sun, Y., Luo, Z., Chen, X., Wang, Y., Qi, B., Lin, J., Lin, W.-W., Sun, C., Zhou, Y., et al. (2022). Exercise Modifies the Transcriptional Regulatory Features of Monocytes in Alzheimer’s Patients: A Multi-Omics Integration Analysis Based on Single Cell Technology. Front. Aging Neurosci. 14, 881488. 10.3389/fnagi.2022.881488.

63. Borggrewe, M., Grit, C., Den Dunnen, W.F.A., Burm, S.M., Bajramovic, J.J., Noelle, R.J., Eggen, B.J.L., and Laman, J.D. (2018). VISTA expression by microglia decreases during inflammation and is differentially regulated in CNS diseases. Glia 66, 2645–2658. 10.1002/glia.23517.

64. Powell, D.R., Blasky, A.J., Britt, S.G., and Artinger, K.B. (2013). Riding the crest of the wave: parallels between the neural crest and cancer in epithelial-to-mesenchymal transition and migration. Wiley Interdiscip. Rev. Syst. Biol. Med. 5, 511–522. 10.1002/wsbm.1224.

65. Stemmler, M.P., Eccles, R.L., Brabletz, S., and Brabletz, T. (2019). Non-redundant functions of EMT transcription factors. Nat. Cell Biol. 21, 102–112. 10.1038/s41556-018-0196-y.

66. Huang, X., and Huang, Y. (2021). Cellsnp-lite: an efficient tool for genotyping single cells. Bioinformatics 37, 4569–4571. 10.1093/bioinformatics/btab358.

67. Huang, Y., McCarthy, D.J., and Stegle, O. (2019). Vireo: Bayesian demultiplexing of pooled single-cell RNA-seq data without genotype reference. Genome Biol. 20, 273. 10.1186/s13059-019-1865-2.

68. Zhu, Y., Wang, L., Yin, Y., and Yang, E. (2017). Systematic analysis of gene expression patterns associated with postmortem interval in human tissues. Sci. Rep. 7, 5435. 10.1038/s41598-017-05882-0.

69. Korsunsky, I., Millard, N., Fan, J., Slowikowski, K., Zhang, F., Wei, K., Baglaenko, Y., Brenner, M., Loh, P., and Raychaudhuri, S. (2019). Fast, sensitive and accurate integration of single-cell data with Harmony. Nat. Methods 16, 1289–1296. 10.1038/s41592-019-0619-0.

70. Finak, G., McDavid, A., Yajima, M., Deng, J., Gersuk, V., Shalek, A.K., Slichter, C.K., Miller, H.W., McElrath, M.J., Prlic, M., et al. (2015). MAST: a flexible statistical framework for assessing transcriptional changes and characterizing heterogeneity in single-cell RNA sequencing data. Genome Biol. 16, 278. 10.1186/s13059-015-0844-5.

71. Bakken, T.E., Jorstad, N.L., Hu, Q., Lake, B.B., Tian, W., Kalmbach, B.E., Crow, M., Hodge, R.D., Krienen, F.M., Sorensen, S.A., et al. (2021). Comparative cellular analysis of motor cortex in human, marmoset and mouse. Nature 598, 111–119. 10.1038/s41586-021-03465-8.

72. Chen, E.Y., Tan, C.M., Kou, Y., Duan, Q., Wang, Z., Meirelles, G.V., Clark, N.R., and Ma’ayan, A. (2013). Enrichr: interactive and collaborative HTML5 gene list enrichment analysis tool. BMC Bioinformatics 14, 128. 10.1186/1471-2105-14-128.

73. Kuleshov, M.V., Jones, M.R., Rouillard, A.D., Fernandez, N.F., Duan, Q., Wang, Z., Koplev, S., Jenkins, S.L., Jagodnik, K.M., Lachmann, A., et al. (2016). Enrichr: a comprehensive gene set enrichment analysis web server 2016 update. Nucleic Acids Res 44, W90–7. 10.1093/nar/gkw377.

74. Xie, Z., Bailey, A., Kuleshov, M.V., Clarke, D.J.B., Evangelista, J.E., Jenkins, S.L., Lachmann, A., Wojciechowicz, M.L., Kropiwnicki, E., Jagodnik, K.M., et al. (2021). Gene Set Knowledge Discovery with Enrichr. Curr. Protoc. 1, e90. 10.1002/cpz1.90.

75. Yu, G., Wang, L.-G., and He, Q.-Y. (2015). ChIPseeker: an R/Bioconductor package for ChIP peak annotation, comparison and visualization. Bioinformatics 31, 2382–2383. 10.1093/bioinformatics/btv145.

76. Castro-Mondragon, J.A., Riudavets-Puig, R., Rauluseviciute, I., Berhanu Lemma, R., Turchi, L., Blanc-Mathieu, R., Lucas, J., Boddie, P., Khan, A., Manosalva Pérez, N., et al. (2022). JASPAR 2022: the 9th release of the open-access database of transcription factor binding profiles. Nucleic Acids Res. 50, D165–D173. 10.1093/nar/gkab1113.

77. Bulik-Sullivan, B.K., Loh, P.R., Finucane, H.K., Ripke, S., Yang, J., Schizophrenia Working Group of the Psychiatric Genomics, C., Patterson, N., Daly, M.J., Price, A.L., and Neale, B.M. (2015). LD Score regression distinguishes confounding from polygenicity in genome-wide association studies. Nat Genet 47, 291–295. 10.1038/ng.3211.

78. Bipolar Disorder and Schizophrenia Working Group of the Psychiatric Genomics Consortium. (2018). Genomic Dissection of Bipolar Disorder and Schizophrenia, Including 28 Subphenotypes. Cell 173, 1705-1715.e16. 10.1016/j.cell.2018.05.046.

79. Landt, S.G., Marinov, G.K., Kundaje, A., Kheradpour, P., Pauli, F., Batzoglou, S., Bernstein, B.E., Bickel, P., Brown, J.B., Cayting, P., et al. (2012). ChIP-seq guidelines and practices of the ENCODE and modENCODE consortia. Genome Res. 22, 1813–1831. 10.1101/gr.136184.111.

80. Jiang, Y., Matevossian, A., Huang, H.-S., Straubhaar, J., and Akbarian, S. (2008). Isolation of neuronal chromatin from brain tissue. BMC Neurosci. 9, 42. 10.1186/1471-2202-9-42.

81. Savic, D., Gertz, J., Jain, P., Cooper, G.M., and Myers, R.M. (2013). Mapping genome-wide transcription factor binding sites in frozen tissues. Epigenetics Chromatin 6, 30. 10.1186/1756-8935-6-30.

82. Reddy, T.E., Pauli, F., Sprouse, R.O., Neff, N.F., Newberry, K.M., Garabedian, M.J., and Myers, R.M. (2009). Genomic determination of the glucocorticoid response reveals unexpected mechanisms of gene regulation. Genome Res. 19, 2163–2171. 10.1101/gr.097022.109.

83. Martin, M. (2011). Cutadapt removes adapter sequences from high-throughput sequencing reads. EMBnet.journal 17, 10–12. 10.14806/ej.17.1.200.

