## Supplementary material for "Single nucleus multiomics identifies ZEB1 and MAFB as candidate regulators of Alzheimer’s disease-specific *cis* regulatory elements": Description of Supplemental Tables

**TableS1_** **TableS1_Donor_Information_Sequencing_Stats.csv**

Donor information associated with DLPFC nuclei used for single nucleus multiomics, related to Figure 1.

All quality control statistics are reported as mean values.

**TableS2_Subcluster_DEGs.csv**

DEGs for one subcluster versus all other subclusters of the same cell type. DEGs were found with MAST with age and sex as covariates. Genes must be expressed in at least 10% of the cells in the subcluster. Significant genes are those with an absolute log2FC>0.5 and an adjusted p-value<0.01

p_val: unadjusted p-value for MAST

avg_log2FC: log fold-chage of the average expression between the two groups. Positive values indicate that the gene is more highly expressed in the subcluster indicated

pct.1: Percentage of cells in the subcluster that express the gene

pct.2: Percentage of cells in other subclusters of the same cell type that express the gene

p_val_adj: Bonferroni adjusted p-value

cluster: cell type and subcluster where the gene is differentially expressed

**TableS3_ADCtrl_DEGs.csv**

DEGs for AD versus control within each cell type. MAST was used with age and sex as covariates. Genes must be expressed in at least 25% of the cells in that cell type. Significant genes have an adjusted p-value less than 0.01 and an absolute log2FC>0.25.

p_val: unadjusted p-value for MAST

avg_log2FC: log fold-chage of the average expression between AD and control. Positive values indicate that the gene is more highly expressed in AD.

pct.1: Percentage of AD cells in the cell type that express the gene

pct.2: Percentage of control cells in the cell type that express the gene

p_val_adj: Bonferroni adjusted p-value

celltype: cell type where the gene is differentially expressed

**TableS4_ADCtrl_DEG_GOenrichment.csv**

GO Biological Process 2021 enrichment for AD v control DEGs split by cell type and effect direction. Significant GO term have adjusted p-value<0.01.

Term: GO Biological Process Term

Overlap: Number of genes that overlap the GO gene list

P.value: unadjusted p-value

Adjusted.P.value: Bonferroni adjusted p-value

Old.P.value: Enrichr old p-value

Old.Adjusted.P.value: Enrichr old adjusted p-value

Odds.Ratio: Odds Ratio for gene set overlap

Combined.Score: ln(p) *z

Genes: Genes that overlap GO gene list

Celltype: Cell type where the genes were differentially expressed

DEGs: direction of log2FC in AD v control

**TableS5_Feature_Linkages.csv**

Merged AD and control feature linkages showing correlation between peak accessibility and gene expression.

Seqnames: Chromosome of linked-peak

Start: start pos of linked-peak

End: end pos of linked-peak

Link: Named Link (gene – peak index #)

Gene: linked-gene

Score.x: Correlation score in AD

Score.y: correlation score in control

Group: AD-only significant in AD; Control-only significant in control; common-significant in both

CT: all cell types that the peak was called in

Astrocytes: Binary indicating if peak was called in astrocytes

Microglia: Binary indicating if peak was called in microglia

Excitatory: Binary indicating if peak was called in excitatory

Inhibitory: Binary indicating if peak was called in inhibitory

Oligodendrocytes: Binary indicating if peak was called in oligodendrocytes

OPCs: Binary indicating if peak was called in OPCs

Signac.seqnames: chromosome name for link plotting

Signac.start: start for link plotting

signac.endL end for link plotting

k27: Cell types that overlapped H3K27ac of the corresponding cell type from Nott et al. 2019 or Kozlenkov et al., 2018

annotation : ChIPseeker peak annotation

distanceToTSS: distance to closest TSS

ATAC_num: Number of cell types that the peak was called in

K27_num: Number of cell types that H3K27ac peak overlapped

Index: peak index number

Score: Average score if link was common

Bin: Binned score

Qval.x: -log10(adjusted pval) for the correlation in AD

Qval.y: -log10(adjusted pval) for the correlation in control

PLACseq_neun: link overlap with neuron PLACseq from Nott et al. 2019

PLACseq_mic: link overlap with microglia PLACseq from Nott et al. 2019

PLACseq_olig: link overlap with oligodendrocyte PLACseq from Nott et al. 2019

CooperMPRA: Linked-peak overlap with MPRA from Cooper et al. 2022.

Weiss_NPC: Linked-peak overlap with MPRA from neural precursor cells (NPCs) from Weiss et al. 2021

Weiss_ESC: Linked-peak overlap with MPRA from ESCs from Weiss et al. 2021

Uebbing_MPRA: Linked-peak overlap with MPRA from NPCs from Uebbing et al. 2021

SuRE_K562: Linked-peak overlap with K562 SuREseq from van Arensbergen et al. 2019

SuRE_HePG2: Linked-peak overlap with HepG2 SuREseq from van Arensbergen et al. 2019

GTEx_FC: link overlap with Frontal Cortex GTEx eQTLs

eQTL_Bryois: link overlap with eQTLs from Bryois et al. 2022

Hu_HiC: link overlap with HiC loops from Hu et al. 2021

**TableS6_Trios.csv**

Trios that showed significant correlation between peak accessibility and gene expression, peak accessibility and TF expression, as well as TF expression and gene expression.

Column names repeated from TableS5 are the same.

GRN: peak-TF motif-gene trios

Motif: motif called in the linked-peak

TF: TF for the motif called in the peak

Motif_score: -log10(pval) of motifmatchr motif call

Motif_seqnames: chromosome of motif call

Motif_start: start pos of motif call

Motif_end: end pos of motif call

G.TF_cor.x: Pearson correlation between gene expression and TF expression in AD

G.TF_pval.x: pval for gene-TF correlation in AD

P.TF_cor.x: Correlation between peak and TF in AD

P.TF_cor.x: pval for peak-TF correlation in AD

G.TF_cor.y: Correlation between gene and TF in control

G.TF_pval.y: pval for gene-TF correlation in control

P.TF_cor.y: Correlation between peak and TF in control

P.TF_pval.y: pval for peak-TF correlation in control

AD_sig: Is gene-TF cor significant in AD?

Ctrl_sig: Is gene-TF cor significant in control?

AD_sigP: Is peak-TF cor significant in AD?

Ctrl_sigP: Is peak-TF cor significant in control?

Category: trio is either only significant in AD (“AD-only”), only significant in control (“Ctrl-only”), significant in both AD and control but in opposite directions (“DiffDir”), or significant in both in the same direction (“Sig”)

**TableS7_MEF2C_GOenrichment.csv**

GO Biological Process 2021 enrichment for targets of MEF2C from trios. Enrichments are split into MEF2C gene targets where the linked-peak was called in neurons and MEF2C gene targets where the linked-peak was called in microglia.

**TableS8_sLDSC_analysis.csv**

GWAS datasets used for sLDSC analysis.

**TableS9_LuciferaseReferenceElements.xlsx**

Genomic location of luciferase elements, genes linked to peaks that are represented by luciferase elements, and individual replicate data for luciferase assays where reference elements are compared to control with only minP activity.

Chromosome: chromosome number

Start Coordinate (hg38): position of first base in designed element

End Coordinate (hg38): position of last base in designed element

Linked Gene: gene(s) linked to peak luciferase elements were designed within

Cell Line Tested: all luciferase assays were performed in either HEK293 or 293FT cells

Mean Diff: Difference between the average of element replicates and control replicates

Unadjusted p-value: calculated by ANOVA with Fisher’s LSD comparing elements to control

Q-value (FDR <0.05): calculated by ANOVA with post hoc FDR of 0.05 comparing elements to control

Rep1-18: Each column is a biological replicate. Biological replicates are the median of 4 technical luminescence values per experiment

**TableS10_LuciferaseSNPelements.xlsx**

Genomic location of luciferase elements, genes linked to peaks that are represented by luciferase elements, SNPs tested, and individual replicate data for luciferase assays performed in HEK293 cells where alternate elements are compared to corresponding reference elements.

Chromosome: chromosome number

Start Coordinate (hg38): position of first base in designed element

End Coordinate (hg38): position of last base in designed element

rsID: rsID for SNP represented in each alternate element. “na” indicates a reference element

Linked Gene: gene(s) linked to peak luciferase elements were designed within

Mean Diff: Difference between the average of alternate element replicates and reference element replicates

Unadjusted p-value: calculated by ANOVA with FIsher’s LSD comparing alternate elements to reference elements

Q-value (FDR <0.05): calculated by ANOVA with post hoc FDR of 0.05 comparing alternate elements to reference elements

Rep1-18: Each column is a biological replicate. Biological replicates are the median of 4 technical luminescence values per experiment

**TableS11_Vireo_demux_stats.csv**

SampleID assignment for 10X Genomics Multiome lanes demultiplexed with cellSNP/vireo where 2 samples were pooled on the same lane.

Sample ID: 10X Genomics library id

Male ID: sample name for male sample in the pool

Female ID: sample name for female sample in the pool

Male cell count: number of cells assigned to the male sample

Female cell count: number of cells assigned to the female sample

Doublets: Number of cells determined to be doublets by vireo

Unassigned: Number of cells with too few snps to assign sample origin

% Assigned: The proportion of cells assigned to a sample
